# Smaller total and subregional cerebellar volumes in posttraumatic stress disorder: a mega-analysis by the ENIGMA-PGC PTSD workgroup

**DOI:** 10.1101/2022.10.13.512111

**Authors:** Ashley A. Huggins, C. Lexi Baird, Melvin Briggs, Sarah Laskowitz, Samar Foudra, Courtney Haswell, Delin Sun, Lauren E. Salminen, Neda Jahanshad, Sophia I. Thomopoulos, Dick J. Veltman, Jessie L. Frijling, Miranda Olff, Mirjam van Zuiden, Saskia B.J. Koch, Laura Nawjin, Li Wang, Ye Zhu, Gen Li, Dan J. Stein, Johnathan Ipser, Soraya Seedat, Stefan du Plessis, Leigh L. van den Heuvel, Benjamin Suarez-Jimenez, Xi Zhu, Yoojean Kim, Xiaofu He, Sigal Zilcha-Mano, Amit Lazarov, Yuval Neria, Jennifer S. Stevens, Kerry J. Ressler, Tanja Jovanovic, Sanne JH van Rooij, Negar Fani, Anna R. Hudson, Sven C. Mueller, Anika Sierk, Antje Manthey, Henrik Walter, Judith K. Daniels, Christian Schmahl, Julia I. Herzog, Pavel Říha, Ivan Rektor, Lauren A.M. Lebois, Milissa L. Kaufman, Elizabeth A. Olson, Justin T. Baker, Isabelle M. Rosso, Anthony P. King, Isreal Liberzon, Mike Angstadt, Nicholas D. Davenport, Scott R. Sponheim, Seth G. Disner, Thomas Straube, David Hofmann, Rongfeng Qi, Guang Ming Lu, Lee A. Baugh, Gina L. Forster, Raluca M. Simons, Jeffrey S. Simons, Vincent A. Magnotta, Kelene A. Fercho, Adi Maron-Katz, Amit Etkin, Andrew S. Cotton, Erin N. O’Leary, Hong Xie, Xin Wang, Yann Quidé, Wissam El-Hage, Shmuel Lissek, Hannah Berg, Steven Bruce, Josh Cisler, Marisa Ross, Ryan J. Herringa, Daniel W. Grupe, Jack B. Nitschke, Richard J. Davidson, Christine Larson, Terri A. deRoon-Cassini, Carissa W. Tomas, Jacklynn M. Fitzgerald, Jennifer Urbano Blackford, Bunmi O. Olatunji, William S. Kremen, Michael J. Lyons, Carol E. Franz, Evan M. Gordon, Geoffrey May, Steven M. Nelson, Chadi G. Abdallah, Ifat Levy, Ilan Harpaz-Rotem, John H. Krystal, Emily L. Dennis, David F. Tate, David X. Cifu, William C. Walker, Elizabeth A. Wilde, Ian H. Harding, Rebecca Kerestes, Paul M. Thompson, Rajendra Morey

## Abstract

**Background:** The cerebellum critically contributes to higher-order cognitive and emotional functions such fear learning and memory. Prior research on cerebellar volume in PTSD is scant and has neglected neuroanatomical subdivisions of the cerebellum that differentially map on to motor, cognitive, and affective functions.

**Methods:** We quantified cerebellar lobule volumes using structural magnetic resonance imaging in 4,215 adults (PTSD n= 1640; Control n=2575) across 40 sites from the from the ENIGMA-PGC PTSD working group. Using a new state-of-the-art deep-learning based approach for automatic cerebellar parcellation, we obtained volumetric estimates for the total cerebellum and 28 subregions. Linear mixed effects models controlling for age, gender, intracranial volume, and site were used to compare cerebellum total and subregional volume in PTSD compared to healthy controls. The Benjamini-Hochberg procedure was used to control the false discovery rate (*p*_-FDR_ < .05).

**Results:** PTSD was associated with significant grey and white matter reductions of the cerebellum. Compared to controls, people with PTSD demonstrated smaller total cerebellum volume. In addition, people with PTSD showed reduced volume in subregions primarily within the posterior lobe (lobule VIIB, crus II), but also the vermis (VI, VIII), flocculonodular lobe (lobule X), and cerebellar white matter (all *p*_-FDR_ < 0.05). Effects of PTSD on volume were consistent, and generally more robust, when examining symptom severity rather than diagnostic status.

**Conclusions:** These findings implicate regionally specific cerebellar volumetric differences in the pathophysiology of PTSD. The cerebellum appears to play an important role in high-order cognitive and emotional processes, far beyond its historical association with vestibulomotor function. Further examination of the cerebellum in trauma-related psychopathology will help to clarify how cerebellar structure and function may disrupt cognitive and affective processes at the center of translational models for PTSD.

## Introduction

Exposure to trauma is common, and nearly 10% of trauma survivors develop chronic symptoms of posttraumatic stress disorder (PTSD; (1)), a debilitating psychiatric condition characterized by a constellation of symptoms including intrusive memories, avoidance, hypervigilance, and negative changes in mood and cognition (2). An extensive body of research has illuminated key brain regions that differentiate PTSD patients from trauma-exposed controls (3–5). Notably, PTSD has been consistently linked to smaller volume of brain regions including the hippocampus (6–9), ventromedial prefrontal cortex (vmPFC; (10–12)), amygdala (13–15), insula (16–18), and anterior cingulate cortex (ACC; (9, 19, 20)). These regions are part of a critical neural circuit supporting diverse cognitive and affective functions that are disrupted in PTSD, including threat processing, emotion regulation, and emotional memory(21, 22).

Relatively little attention has been paid to areas of the brain outside these canonical regions. Notably, research emerging over the past three decades clearly demonstrates that the cerebellum contributes immensely to higher-order cognition and emotion (23–25). Historically known for its central role in the vestibulomotor system (26), the human cerebellum has rapidly (and disproportionately) evolved over time (27–29). Despite being approximately 10% of the brain’s overall size (30), the cerebellum houses the vast majority of the brain’s total neurons (31) and occupies nearly 80% of the neocortical surface area (29). The cerebellum shares rich anatomical connections with much of the brain, including with prefrontal and limbic areas (27, 32–34), strongly suggesting that it participates in processes beyond motor coordination that may be highly relevant to PTSD. Moreover, the cerebellum’s widespread connectivity with stress-related regions (such as with the amygdala, hippocampus, and periaqueductal gray) may make it especially vulnerable to traumatic stress, potentially leading to the development of PTSD symptoms by disrupting typical brain-mediated stress responses via cerebro-cerebellar circuits (35, 36). Recent studies have also demonstrated that the cerebellum is involved in fear learning and memory (37–40); considering PTSD is characterized by aberrancies in threat detection and processing (41, 42), this accumulating evidence makes a compelling case that the cerebellum is involved in the pathophysiology of PTSD.

A growing body of structural and functional magnetic resonance imaging studies provide evidence of altered cerebellar volume and function in PTSD (37). Specifically, smaller cerebellar volume has been observed in both adult (43, 44) and pediatric (45, 46) PTSD samples. PTSD has also been linked to disrupted functional connectivity between the cerebellum and key cognitive and affective regions, including the amygdala (47). Although meta-analytic work has suggested cerebellar activation differentiates PTSD patients from healthy controls (48–50), other studies have failed to observe any cerebellar volumetric differences related to PTSD (51–53), necessitating additional studies to resolve these discrepant findings. Collectively, these results highlight the importance of incorporating the cerebellum into well-established translational models of PTSD.

Prior research on cerebellar volume in PTSD has been limited by largely neglecting to consider important neuroanatomical subdivisions of the cerebellum that differentially map onto motor, cognitive, and affective functions. Gross anatomy delineates two major fissures dividing the cerebellum into three anatomical divisions: the anterior (lobules I-V), posterior (lobules VI-IX), and flocculonodular (lobule X) lobes (54). The anterior lobe receives spinal afferents via spinocerebellar tracts and shares reciprocal connections with motor cortices to help support motor movements, gait, and equilibrium (55). By contrast, extensive non-motor functions have been identified within the evolutionarily newer posterior cerebellum (56), which lacks spinal cord inputs and has connections with cortical areas integral to higher order processes, including the prefrontal cortex and cingulate gyrus (57, 58). Activation within the posterior lobe has been observed during language and verbal working memory (lobule VI, crus I), spatial processing (lobule VI), and executive function (lobule VI and VIIB, crus I) tasks (24, 56, 59). Aversive stimulus processing, such as noxious heat and unpleasant images, also appears to involve the posterior cerebellum (lobules VI and VIIB and crus I), implicating these regions in defensive responding (60). The vermis - the medial cortico-nuclear column connecting the left and right cerebellar hemispheres - is considered an extension of the Papez emotion circuit (61) and is activated during affective processing (23, 25, 62). Vermal lobules also interact with other regions critical for emotional associative learning including the amygdala, hypothalamus, and periaqueductal gray (23, 63, 64). Taken together, these careful studies on functional topography have identified three broad subdivisions of the cerebellum comprising sensorimotor, cognitive, and limbic areas (24).

As a heterogenous disorder linked to dysfunction within multiple cerebellum-supported processes, it is unclear whether structural differences in the cerebellum in PTSD are global or may be localized to specific subregions. Indeed, prior work has identified differences in cerebellar volume and function distributed across the cerebellum, including within the vermis (43, 46), crus (44, 65), and lobules VI and VII (66–68). Yet, these diffuse subregional findings are often not replicated, contributing to a lack of consensus regarding the cerebellum’s role in PTSD. Importantly, better understanding the relevance of cerebellar structure in the pathophysiology of PTSD may help elucidate potential mechanisms that perpetuate chronic symptoms of PTSD and aid in our ability to develop targeted, effective interventions.

To this end, the present study employed a mega-analysis of total and subregional cerebellar volumes in a large, multi-cohort dataset from the Enhancing NeuroImaging Genetics through Meta-Analysis (ENIGMA)-Psychiatric Genomics Consortium (PGC) PTSD workgroup. By contrast with a meta-analysis, a mega-analysis centralizes and pools data from multiple sites and fits statistical models to the aggregated data while adjusting for site effects. We used a novel, standardized ENIGMA cerebellum parcellation protocol (Kerestes et al., 2022) to quantify cerebellar lobule volumes using structural MRI data from 4,215 adults with (n=1,640) and without (n=2,575) PTSD. We examined the effects of PTSD on cerebellar volumes, adjusting for age, gender, and total intracranial volume. Based on prior work (43–46), we hypothesized that PTSD would be associated with smaller total cerebellum volume. Considering functional topography indicates the ‘limbic’ and ‘cognitive’ cerebellum localize to the vermis and posterior lobes, respectively, we hypothesized PTSD would be associated with smaller volumes within these two anatomical divisions (23–25).

## Methods and Materials

### Sample

Clinical, demographic, and neuroimaging data from the ENIGMA-PGC PTSD working group included in the current study are presented in Table 1. MRI scans from 4,215 subjects, including 1640 PTSD patients and 2,575 healthy controls (trauma-exposed or naïve), were automatically segmented into cerebellar subregions. All study procedures were approved by local institutional review boards (IRB), and participants provided written informed consent. The present analyses were granted exempt status by the Duke University Health System IRB.

**Table 1.** Sample characteristics by site.

| Site | N | Age | Gender |  | Diagnosis |  | Diagnostic Tool | Sample | PTSD Severity | Severity Tool |
| --- | --- | --- | --- | --- | --- | --- | --- | --- | --- | --- |
|  |  | *M*<br>(*SD*) | *F* | *M* | *PTSD* | *Ctrl* |  |  | *M* %<br>(*SD*) |  |
| ADNI DoD | 103 | 68.53<br>(4.05) | 2 | 101 | 43 | 60 | CAPS-IV | Military | 21.00<br>(20.71) | CAPS-IV |
| Amsterdam AMC | 73 | 40.01<br>(9.97) | 34 | 39 | 36 | 37 | CAPS-IV | Police | 26.37<br>(24.57) | CAPS-IV |
| Beijing | 87 | 48.49<br>(10.29) | 53 | 34 | 41 | 46 | PCL-5 | Civilian | 35.52<br>(20.00) | PCL-5 |
| Cape Town | 106 | 26.78<br>(6.39) | 106 | 0 | 6 | 100 | CAPS-IV | Civilian | 46.14<br>(21.74) | CAPS-IV |
| Columbia | 151 | 34.96<br>(10.65) | 90 | 61 | 72 | 79 | CAPS-IV,<br>SCID | Civilian | 34.07<br>(26.55) | CAPS-IV, CAPS-5 |
| Duke | 376 | 39.24<br>(10.02) | 72 | 304 | 111 | 265 | CAPS-IV,<br>CAPS-5, SCID | Military | 20.04<br>(23.44) | CAPS-IV, CAPS-5, DTS |
| Emory GTP | 59 | 40.24<br>(11.97) | 59 | 0 | 13 | 46 | CAPS-IV | Civilian | 21.04<br>(16.31) | CAPS-IV |
| Ghent | 65 | 37.15<br>(12.22) | 65 | 0 | 8 | 57 | MINI | Civilian | --- | --- |
| Groningen | 37 | 38.81<br>(9.46) | 37 | 0 | 37 | 0 | CAPS-IV | Civilian | 49.24<br>(9.63) | CAPS-IV |
| LIMBIC-CENC | 1045 | 40.10<br>(9.84) | 144 | 901 | 354 | 691 | PCL-5 | Military | 32.12<br>(23.73) | PCL-5 |
| Mannheim | 40 | 35.83<br>(11.50) | 40 | 0 | 40 | 0 | SCID | Civilian | 53.95<br>(22.26) | DTS |
| Masaryk | 269 | 51.72<br>(18.67) | 166 | 103 | 109 | 160 | PCL-C | Civilian | 34.59<br>(12.00) | PCL-C |
| McLean 1 | 78 | 34.56<br>(12.49) | 78 | 0 | 51 | 27 | CAPS-5 | Civilian | 42.85<br>(31.99) | CAPS-5 |
| McLean 2 | 94 | 34.15<br>(8.89) | 52 | 42 | 21 | 73 | CAPS-IV | Civilian | --- | CAPS-IV |
| Michigan | 62 | 30.42<br>(7.71) | 0 | 62 | 40 | 22 | CAPS-IV | Military,<br>Civilian | 35.92<br>(25.10) | CAPS-IV |
| Milwaukee | 70 | 32.48<br>(10.05) | 35 | 35 | 19 | 51 | CAPS-5 | Civilian | 17.45<br>(15.34) | CAPS-5 |
| Minnesota | 62 | 42.85<br>(9.51) | 5 | 57 | 12 | 50 | CAPS-IV | Military | 13.54<br>(14.39) | CAPS-IV |
| Missouri | 64 | 32.02<br>(9.73) | 64 | 0 | 57 | 7 | CAPS-IV | Civilian | --- | --- |
| Münster | 43 | 26.41<br>(6.85) | 38 | 5 | 19 | 24 | SCID | Civilian | --- | --- |
| Nanjing | 132 | 57.23<br>(5.94) | 73 | 59 | 48 | 84 | SCID | Civilian | 20.83<br>(13.44) | CAPS-IV |
| South Dakota | 114 | 29.29<br>(10.44) | 21 | 93 | 71 | 43 | PCL-C, PCL-M | Military | 44.67<br>(18.56) | PCL-C, PCL-M |
| Stanford | 146 | 33.59<br>(10.44) | 67 | 78 | 73 | 73 | CAPS-IV | Military,<br>Civilian | 26.57<br>(23.64) | CAPS-IV |
| Toledo | 77 | 35.38<br>(11.40) | 35 | 42 | 15 | 62 | CAPS-IV | Military,<br>Civilian | 17.49<br>(18.42) | CAPS-IV |
| Tours | 39 | 28.23<br>(9.88) | 39 | 0 | 9 | 30 | CAPS-IV | Civilian | 31.11<br>(14.41) | CAPS-IV |
| Wisconsin 1 | 104 | 33.00<br>(8.25) | 104 | 0 | 83 | 21 | CAPS-5, SCID | Civilian | 47.26<br>(24.62) | PCL-5, PCL-C |
| Wisconsin 2 | 24 | 29.96<br>(5.52) | 3 | 21 | 12 | 12 | CAPS-IV | Military | 25.77<br>(24.68) | CAPS-IV |
| VA Minneapolis | 241 | 32.64<br>(7.80) | 13 | 226 | 91 | 150 | CAPS-IV | Military | 29.40<br>(19.16) | CAPS-IV |
| VA Waco | 91 | 39.75<br>(11.02) | 11 | 80 | 63 | 29 | PCL-5 | Military | 54.10<br>(26.21) | PCL-5 |
| VA West Haven | 55 | 33.55<br>(8.77) | 6 | 49 | 32 | 23 | CAPS-IV | Military | 34.16<br>(21.94) | CAPS-IV |
| Vanderbilt | 46 | 31.24<br>(4.64) | 9 | 37 | 12 | 34 | CAPS-5 | Military | 10.76<br>(14.49) | CAPS-5 |
| VETSA | 190 | 61.79<br>(2.66) | 0 | 190 | 19 | 171 | PCL-C | Military | 28.89<br>(11.24) | PCL-C |
| Yale | 69 | 29.61<br>(7.65) | 11 | 58 | 31 | 23 | CAPS-IV | Military | 20.69<br>(20.97) | CAPS-IV |
| **Overall** | **4215** | **40.08**<br>**(13.72)** | **1532** | **2677** | **1640** | **2575** | **---** | **---** | **30.53**<br>**(23.12)** | **---** |

*Note:* CAPS-IV, Clinician Administered PTSD Scale for DSM-IV; CAPS-5, Clinician Administered PTSD Scale for DSM-5; DTS, Davidson Trauma Scale for DSM-IV; MINI, Mini Neuropsychiatric Interview; PCL-C, PTSD Checklist-Civilian Version; PCL-M, PTSD Checklist-Military Version; PCL-5, PTSD Checklist for DSM-5; SCID, Structured Clinical Interview for DSM

### Image acquisition and processing

Whole-brain T1-weighted anatomical MR images were collected from each participant. Acquisition parameters for each cohort are detailed in Supplementary Table S2. Segmentation and quality control procedures were performed at Duke University. A subset of the data (n=1,045) from the Long-Term Impact of Military-Relevant Brain Injury Consortium-Chronic Effects of Neurotrauma Consortium (LIMBIC-CENC) were processed at University of Utah. Cerebellar parcellation was carried out using a deep-learning algorithm, Automatic Cerebellum Anatomical Parcellation using U-Net with Locally Constrained Optimization (ACAPULCO) (69). Images were corrected for intensity inhomogeneity using N4, blurred with a 3D Gaussian kernel (SD=3mm), and transformed to MNI template space. ACAPULCO then employed a cascade of two convolutional neural networks to first define a 3D-bounding box around the cerebellum and then divide it into anatomically meaningful regions. This ultimately resulted in volumetric estimates for the total cerebellum and 28 subregions, including the hemispheric anterior (lobules I-III, IV, and V), posterior (lobules VI, VIIB, VIIIA, VIIIB, IX, and crus I-II), and flocculonodular (lobule X) lobes, vermal lobules VI, VII, VIII, IX, and X, and the corpus medullare (the white matter core of the cerebellum). ACAPULCO achieves results comparable to other established cerebellum parcellation protocols (e.g., CERES2), but may perform better for multi-site datasets (69).

Following segmentation, a two-step quality control procedure was employed, consisting of (1) removal of statistical outliers ± 2.689 SD from the site mean, and (2) visual inspection of cerebellar parcels. Each subject’s segmentation was visually inspected and scored by a minimum of two trained raters (AH, SL, MB, LB) on a scale from 1 (good) to 3 (poor/failed). In the event of a discrepancy between raters, the parcellation was examined by a third rater for consensus. Segments were considered individually; therefore, select subregional volumes (e.g., statistical outliers, circumscribed segmentation errors) were excluded, while the remainder of segments were retained for analysis if correct. Subjects who scored 3 were excluded from all analyses. Breakdown of ratings by site are noted in Supplementary Table S3.

**Figure 1.**
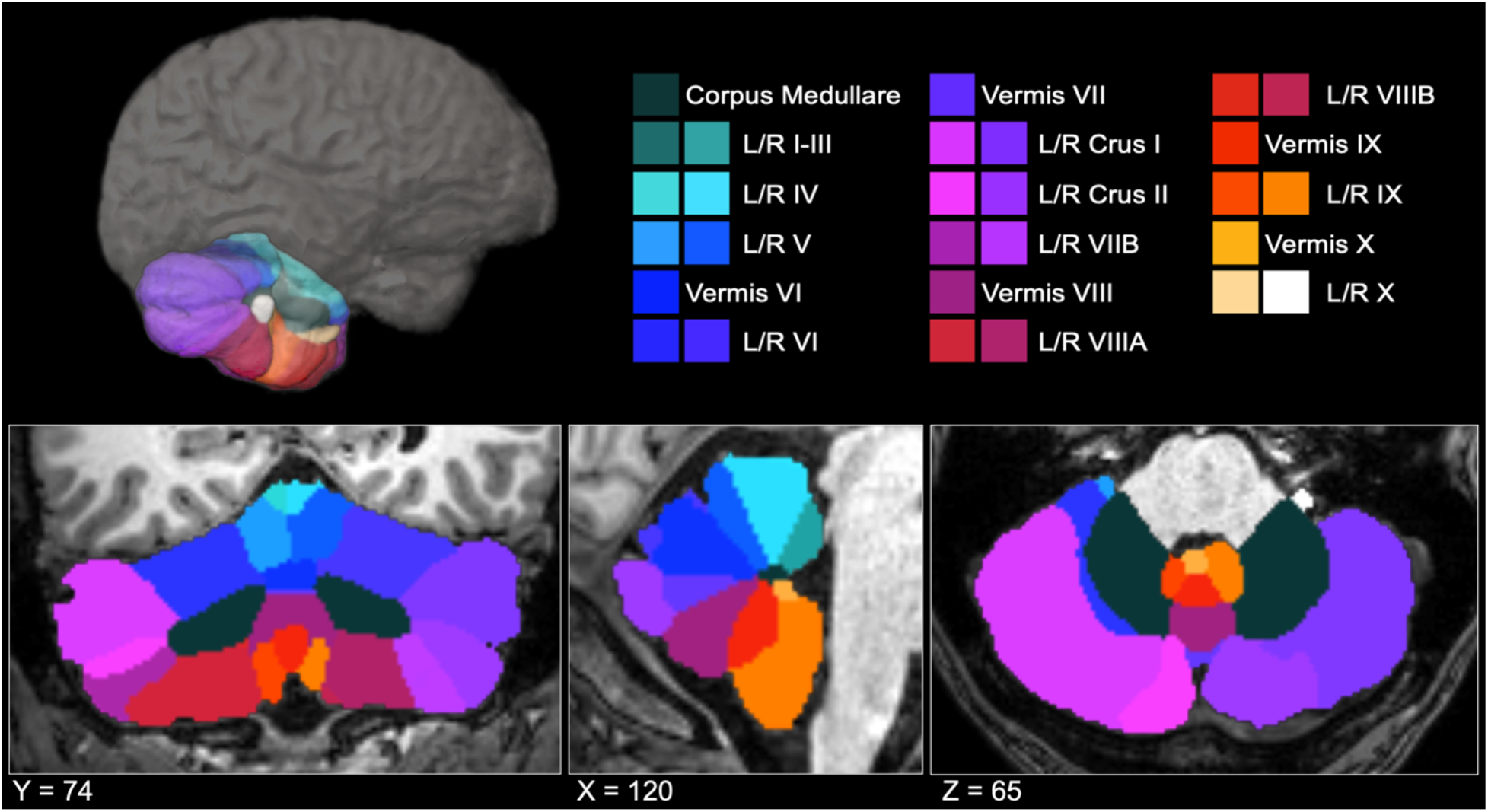
ACAPULCO cerebellum parcellation for a representative subject in three-dimensional (upper left), coronal (left), sagittal (middle), and axial (right) views. L, left; R, right.

### Statistical analysis

To examine whether PTSD diagnosis was associated with volume differences in the grey matter volumes of the whole cerebellum, hemispheric subregions, vermis, and cerebellar white matter, we fit a series of linear mixed effects models were performed. Statistical analyses were conducted using the *lmer* package (70) in R v4.1.3. In each model, age, gender, and total intracranial volume were treated as fixed effects, and site was treated as a random effect. The Benjamini-Hochberg procedure (71) was used to adjust significance values to control the false discovery rate (*p*_-FDR_ < .05). Cohen’s *d* was calculated as a measure of effect size. Models were repeated implementing PTSD severity – rather than diagnosis – as a continuous predictor. Due to site measurement differences, PTSD severity was quantified as a percentage of the total score possible (see Table 1).

Given frequent co-occurrence of PTSD and likely independent effects on cerebellum volume, secondary analyses were conducted to examine the potential effects of depression (72, 73), alcohol use disorder (74, 75), and childhood trauma (76, 77) on cerebellar volumes. For sites with available covariate data (see Supplemental Material), an additional series of linear mixed effects models were conducted, including fixed effects of (1) major depressive disorder diagnosis, (2) alcohol use disorder diagnosis, and (3) total score on the Childhood Trauma Questionnaire (CTQ; (78)).

## Results

### Associations between PTSD diagnosis and cerebellum volumes

Effects of PTSD diagnosis on cerebellum volumes are presented in Table 2. Consistent with hypotheses, after adjusting for age, gender, and total intracranial volume, PTSD diagnosis was associated with significantly smaller total cerebellar volume, *b* = −976.90, *t* = −2.779, *p*_-FDR_ = 0.005. PTSD diagnosis was also associated with smaller volume of the corpus medullare, *b* = −157.47, *t* = −2.234, *p*_-FDR_ = 0.026.

**Table 2:**
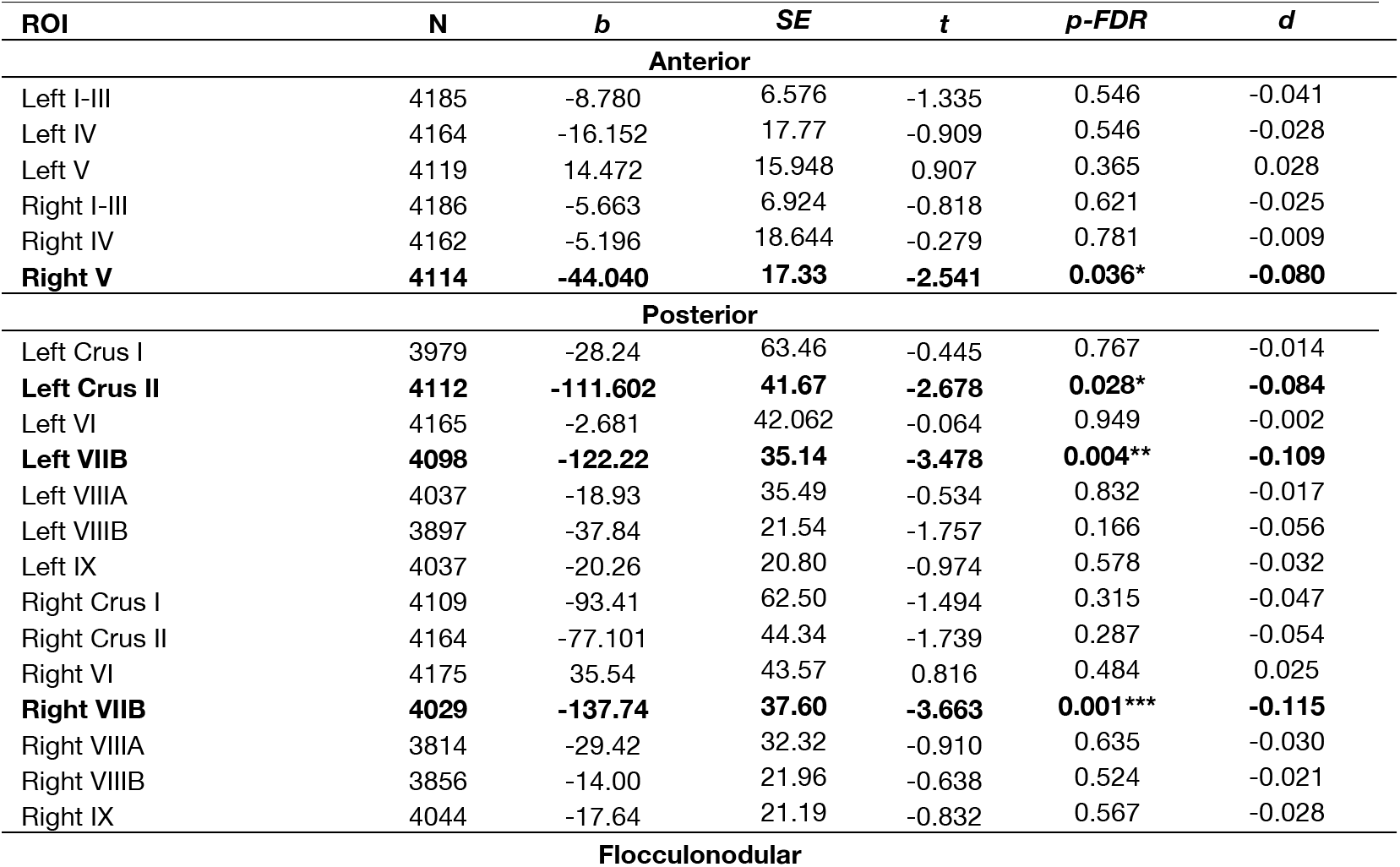
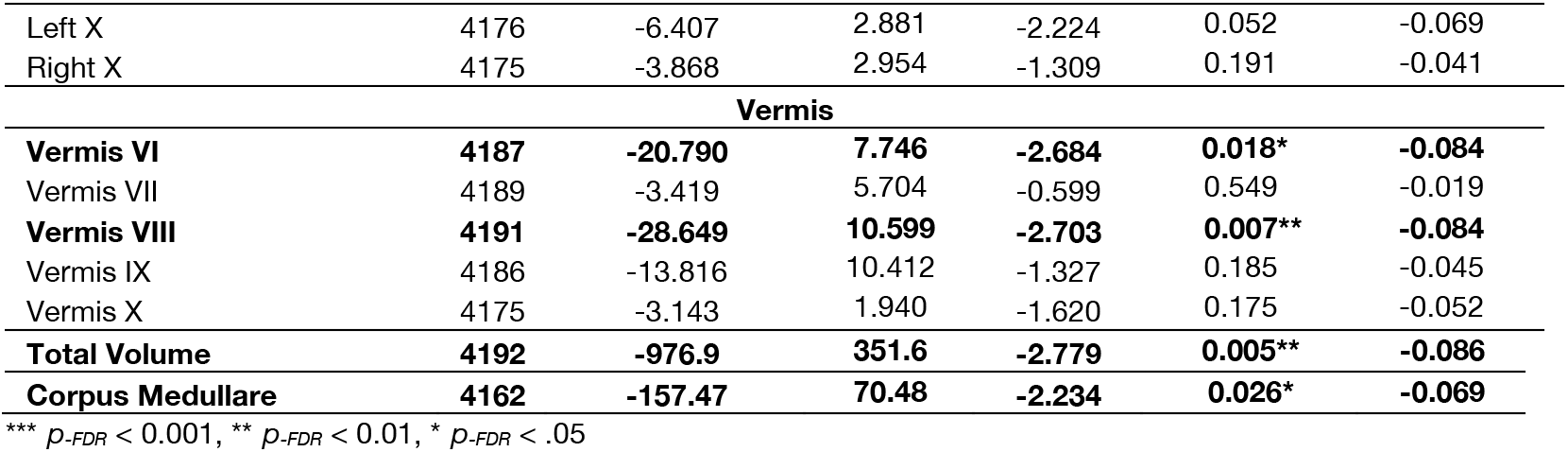
Effects of PTSD diagnosis on cerebellum volume.

Within the anterior cerebellum (lobules I-V), PTSD diagnosis was associated with smaller volume of right lobule V, *b* = −44.040, *t* = −2.541, *p*_-FDR_ = 0.036.

Within the posterior cerebellum (crus, lobules VI-IX), PTSD diagnosis was associated with smaller volume of left crus II, *b* = −111.602, *t* = −2.678, *p*_-FDR_ = 0.028, left lobule VIIB, *b* = −122.22, *t* = −3.478, *p*_-FDR_ =.004, and right lobule VIIB, *b* = −137.74, *t* = −3.663, *p*_-FDR_ = 0.001.

No significant effects of PTSD diagnosis were observed on volumes within the flocculonodular lobe (lobule X). There was an effect of PTSD on left lobule X volume, but this did not survive multiple comparisons corrections (*p*_-FDR_ = 0.052).

There was a significant effect of PTSD diagnosis on volumes of vermal lobules VI, *b* = −20.790, *t* = −2.684, *p*_-FDR_ = 0.018, and VIII, *b* = −28.649, *t* = −2.703, *p*_-FDR_ = 0.007. There were no other significant effects of PTSD within the vermis.

### PTSD severity

When examining PTSD symptom severity (rather than diagnostic status), results were similar, if generally more robust (see Table 3). Specifically, PTSD symptom severity was associated with significantly smaller total cerebellum volume, *b* = −682.00, *t* = −3.688, *p*_-FDR_ = 0.0002, and corpus medullare volumes, *b* = −112.75, *t* = −3.030, *p*_-FDR_ = 0.0002. Effects were consistent across the posterior cerebellum and vermis, with significant effects of PTSD symptom severity on volumes of left crus II, *b* = −64.30, *t* = −2.943 *p*_-FDR_ = 0.011, left lobule VIIB, *b* = −64.83, *t* = −3.529, *p*_-FDR_ = 0.003, right lobule VIIB, *b* = −77.51, *t* = −3.903, *p*_-FDR_ = 0.0007, and vermal lobules VI, *b* = −14.464, *t* = −3.554, *p*_-FDR_ = 0.002, and VIII, *b* = −17.150, *t* = −3.061, *p*_-FDR_ = 0.006.

**Table 3:**
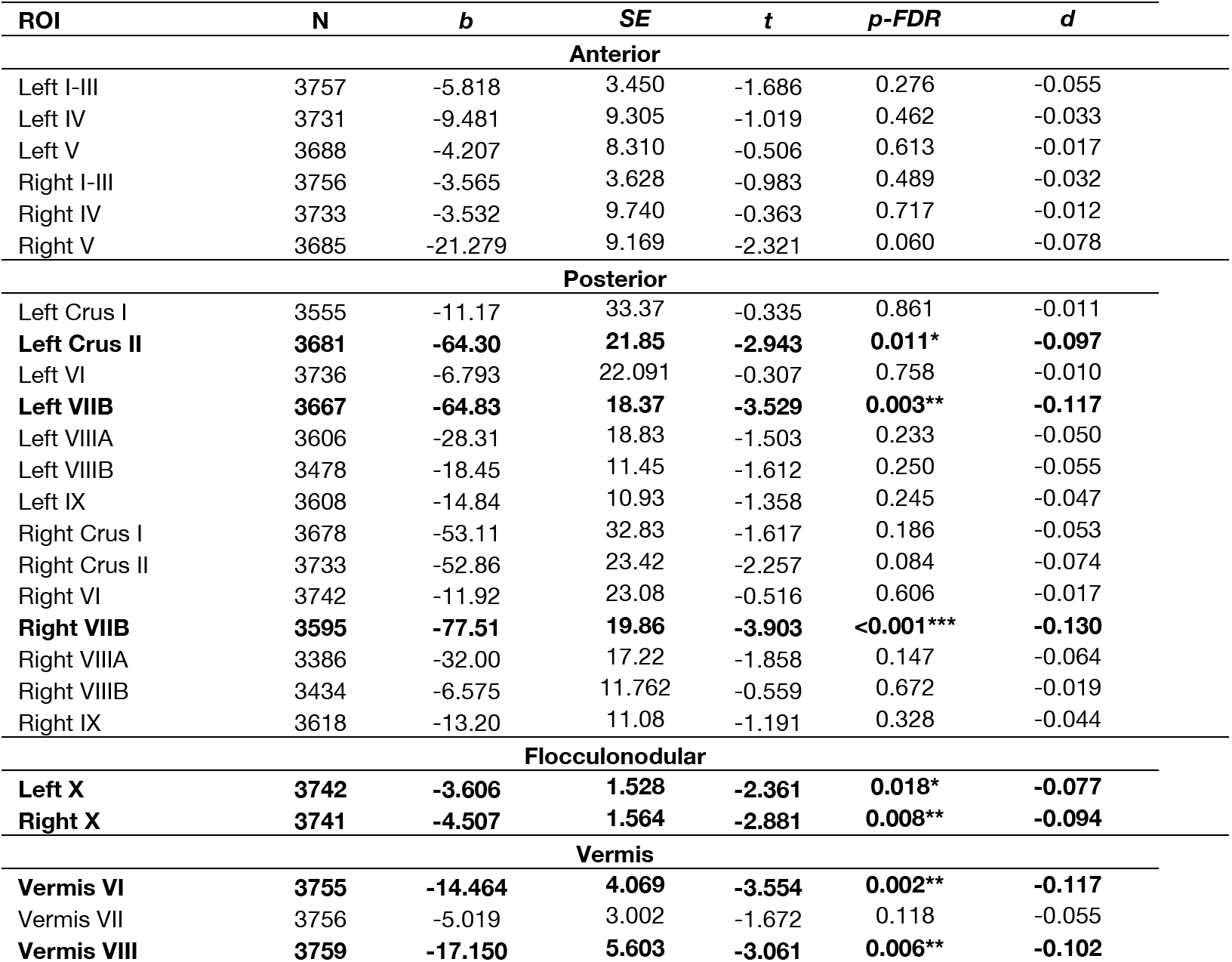
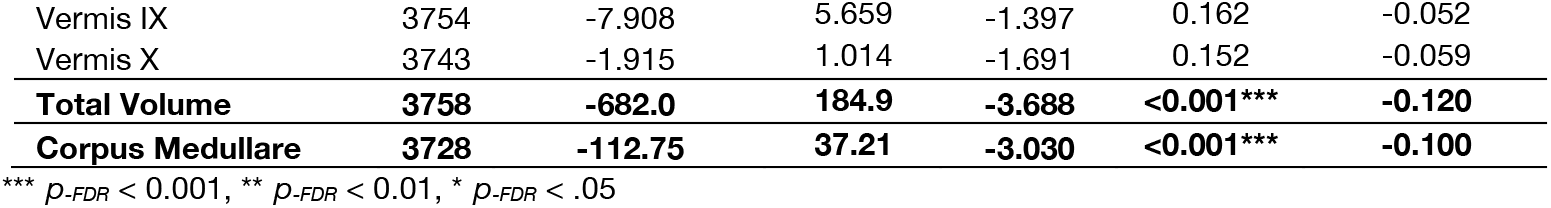
Effects of PTSD severity on cerebellar volumes.

By contrast, the significant effect of PTSD on volume of right lobule V was no longer significant when examining symptom severity instead of diagnosis (*p*_-FDR_ = 0.060). Additionally, PTSD symptom severity was associated with significantly smaller volume of the flocculonodular cerebellum, with effects observed in both hemispheres of lobule X (left: *b* = −3.606, *t* = −2.361, *p*_-FDR_ = 0.018; right: *b* = −4.507, *t* = −2.881, *p*_-FDR_ = 0.008).

### Potential confounding variables

When including covariates assessing depression, alcohol use, and childhood trauma, effects of PTSD on cerebellar volumes were somewhat diminished (See Supplemental Material). Yet, detecting significant effects in these additional analyses presented a challenge to statistical power. There was high collinearity between PTSD and covariates, and - in the case of alcohol use disorder and childhood trauma severity - substantially reduced sample size because not all sites reported these variables. In cases where the effect of PTSD diagnosis was non-significant upon inclusion of covariates, we followed up by testing whether depression, alcohol use, or childhood trauma predicted cerebellar volumes on their own; in *no* instance were covariates found to independently predict cerebellar volumes when PTSD status was excluded from the model, demonstrating that our initial findings were specific to PTSD.

Depression status was available for the majority of subjects (n=3978). When adjusting for major depressive disorder diagnosis, PTSD diagnosis remained significantly associated with smaller volume of both left and right lobule VIIB, and vermis VI. While initially significant, effects of PTSD diagnosis on right lobule V (*p*_-FDR_ = 0.096) and left crus II (*p*_-FDR_ = 0.133) volumes did not survive correction for multiple comparisons. PTSD symptom severity was associated with smaller total cerebellum and vermis VIII volumes. Uniquely, depression diagnosis was associated with smaller volume of right lobule X, *b* = −8.282, *t* = −2.356, *p*_-FDR_ = 0.038.

When adjusting for alcohol use disorder, PTSD was associated with significantly smaller volume of vermal lobule VI. Effects of PTSD diagnosis (*p*_-FDR_ = 0.151) and symptom severity (*p*_-FDR_ = 0.087) on total cerebellar volume did not reach significance when including alcohol use disorder in the model. Including CTQ severity as a covariate resulted in null effects of PTSD diagnosis; significant effects in left lobule VIIB (*p*_-FDR_ = 0.133) and vermal lobule VI (*p*_-FDR_ = 0.075) were no longer significant after correction for multiple comparisons. PTSD symptom severity, however, was significantly associated with vermal lobule VI (*p*_-FDR_ = 0.022) and total cerebellar (*p*_-FDR_ = 0.036) volume after adjusting for childhood trauma.

## Discussion

Leveraging an international, multisite dataset from ENIGMA-PGC PTSD, we conducted a mega-analysis of total and subregional cerebellar volume in PTSD. Consistent with hypotheses based on published work (43–46), PTSD was associated with smaller total cerebellar volume. We found subregional specificity linking PTSD to smaller volumes in the posterior cerebellum, vermis, and flocculonodular cerebellum. Effects of PTSD on cerebellum volume were consistent (and generally more robust) when examining symptom severity rather than diagnostic status. Overall, these findings contribute to an emerging literature that underscores the relevance of cerebellar structure in the pathophysiology of PTSD. Although the appreciation of the cerebellum for its contributions to cognitive and affective function is relatively recent, the current results bolster a growing literature confirming the cerebellum is not exclusively devoted to motor function and may, in fact, have unique relevance to psychiatric conditions including PTSD (34, 37, 79).

Multiple neuroimaging studies have suggested that altered structure and function of the posterior cerebellum may be a neural correlate of PTSD. For instance, structural differences in lobules VIIB, VIIIA, and VIIIB were found in combat-exposed veterans with PTSD (68). Functionally, PTSD has been linked to increased activation during attentional and emotional tasks (66, 67) and decreased resting-state amplitude of low-frequency fluctuation (80) in lobule VI. In a sample of sexual assault survivors, PTSD severity was negatively associated with activation in lobules VI, VIII, IX, and crus I during the performance of an emotional go/no-go task, and positively associated with activation in left cerebellar lobules VII-IX and crus I-II when retrieving positive memory during a mental imagery task (81). PTSD has also been linked to decreased global connectivity within the posterior cerebellum during symptom provocation (82). As the most phylogenetically recent part of the cerebellum (27), the posterior lobe is intricately linked with paralimbic and association cortical areas and plays an integral role in the integration of perception, emotion, and behavior (24, 25). Accordingly, the posterior cerebellum contributes to the salience network (lobules VI and VII; (23, 83)) and diverse cognitive-affective processes including working memory, attentional allocation, and associative learning (24, 84). In the context of the current findings, smaller volume of lobule VIIB and crus II may be implicated in the pathophysiology of PTSD, perhaps mapping directly onto symptoms such as hypervigilance and concentration difficulties.

In the present study, PTSD was also associated with smaller volume of vermal lobules VI and VIII. The cerebellar vermis is considered part of the ‘limbic’ cerebellum and appears to play a key role in emotional processing, learning, and memory (23, 25, 62). Prior work has demonstrated that PTSD is associated with smaller volume (43, 46) and increased signal variability (85) of the vermis. Importantly, structural abnormalities in the vermis may provide increased spatial specificity within existing translational models of PTSD, as converging evidence from both animals and human subjects has shown vermal activation is important for both acquisition (86–89) and extinction (90, 91) of conditioned fear. The cerebellar vermis has strong connections to brain regions (including the brainstem, amygdala, and hypothalamus) that regulate critical survival functions (92). The vermis may contribute to fear learning via threat-associated autonomic changes facilitating defensive behavior, such as increases in respiration, heart rate, and blood pressure (88). Animal research highlights mechanistic links between vermal-midbrain connectivity and defensive behavior; in rats, for instance, lesions of the pathway between the periaqueductal gray and vermal lobule VIII provoke fear-evoked freezing behavior (93). Importantly, vermal connectivity is also implicated in clinical human samples, and PTSD is associated with disrupted resting-state functional connectivity from the vermis to amygdala, periaqueductal gray, and ventromedial prefrontal cortex (94).

Curiously, PTSD symptom severity was associated with reduced volume of bilateral lobule X (which comprises the flocculonodular lobe), but its association with PTSD diagnosis was non-significant. The flocculonodular lobe is primarily implicated in ocular tracking and regulation of the vestibular system (95). Yet, when depression diagnosis was added to the model, there was a significant negative effect of depression on right lobule X, whereas effects of PTSD were non-significant. Structural differences in lobule X have previously been observed in major depressive disorder (96), and these differences have been attributed to somatic complaints, such as dizziness, that are frequently endorsed by patients with depression. PTSD and major depressive disorder are highly comorbid (97, 98). Therefore, smaller lobule X volume may be unique to patients with prominent depressive features and/or a more somatic symptom profile.

## Limitations

This is the largest study of cerebellar volumetry in PTSD to date, however, there are several notable limitations. PTSD is a heterogeneous disorder and is highly comorbid with other psychiatric conditions (e.g., depression, substance use disorders) and environmental exposures (e.g., childhood trauma) that are also linked to alterations in cerebellar structure (72, 77, 79). Employing a mega-analysis in a large multi-cohort consortium dataset enabled us to observe small effect sizes of PTSD on cerebellar volume in our primary analyses, but many sites did not provide diagnostic or item-level data for relevant covariates. Consequently, we were unable to investigate effects of relevant covariates at the same scale. Future studies would benefit from investigating unique and shared phenotypes of PTSD and other psychopathology on the cerebellum to disentangle potential dissociable effects and complex interactions more elegantly. It is also critical for future work to examine how the cerebellum may be uniquely implicated in the dissociative subtype of PTSD. Dissociative symptoms in PTSD are linked to alterations within the midbrain that facilitate passive, rather than active, defensive responses (99, 100); observed differences in cerebellar functional activation and connectivity related to the dissociative subtype of PTSD (65, 67, 101, 102) may be mediated by the prominent neural pathways between the cerebellum and midbrain. The current study was also focused solely on cerebellar volumetric differences in PTSD. Multiple studies have observed disrupted cerebellar activity both at rest (44, 65, 102) and during trauma-relevant tasks (47, 67, 81, 103) in patients with PTSD. Future work would benefit from improved localization of both functional and structural changes in the cerebellum that may be present in PTSD. Lastly, the current study is cross-sectional in nature; future longitudinal research will be imperative to better understand whether cerebellum volume confers risk for PTSD or changes as a function of the disorder.

## Conclusion

In a sample of over 4000 individuals from the ENIGMA-PGC PTSD Consortium, cerebellum volume was significantly smaller in patients with PTSD compared to pooled groups of trauma-exposed and trauma naïve controls. Specific subregional volume reductions in the vermis and posterior cerebellum (crus II and lobule VIIB) align with previous work demonstrating their involvement in cognitive and affective functions relevant to PTSD, such as fear learning and regulation. Overall, these findings argue for a critical role of the cerebellum in the pathophysiology of PTSD, bolstering support for the region’s contributions to processes beyond vestibulomotor function.

## Notes

### Competing Interest Statement

The authors have declared no competing interest.

