## Supplemental Material for "Smaller total and subregional cerebellar volumes in posttraumatic stress disorder: a mega-analysis by the ENIGMA-PGC PTSD workgroup"

**Supplementary analyses:** Method and results for linear mixed effects models including additional covariates (depression, alcohol use disorder, childhood trauma)

**Table S1:** Site inclusion and exclusion criteria
**Table S2:** Scan parameters by site
**Table S3:** Quality control ratings
**Table S4:** Depression characteristics by site
**Table S5:** Alcohol use disorder and childhood trauma characteristics by site
**Table S6:** Race and ethnicity by site
**Table S7:** Effects of depression on cerebellum volumes
**Table S8:** Effects of alcohol use disorder on cerebellum volumes
**Table S9:** Effects of childhood trauma on cerebellum volumes

**Supplementary references**

**Supplementary analyses**

**Major depressive disorder (MDD).** To covary for potential effects of depression on total and subregional cerebellar volumes, we performed linear mixed effects models that included fixed effects of age, gender, intracranial volume, PTSD diagnosis, and MDD diagnosis. MDD diagnosis was coded as a binary variable (0=no current MDD, 1=current MDD). Group status was determined by clinical interview (e.g., SCID) that indicated diagnostic criteria were met for current MDD. Cut-off scores were used to derive group status for subjects without diagnosis but with self-report symptom severity scores. The cut-off scores were used as follows: Beck Depression Inventory-II (BDI-II) ≥ 20 (Beck et al., 1996); Beck Depression Inventory-Short Form (BDI-SF) ≥ 14; Center for Epidemiologic Studies Depression Scale (CES-D) ≥ 16 (Smarr & Keefer, 2011); Depression Anxiety Stress Scales-21 (DASS-21) depression subscale ≥ 14 (Lovibond & Lovibond, 1995); Geriatric Depression Scale (GDS) ≥ 10 (Greenberg, 2007); Hamilton Depression Rating Scale (HAM-D) ≥ 17 (Zimmerman et al., 2013); Hospital Anxiety and Depression Scale-Depression (HADS-D) ≥ 8 (Brennan et al., 2010); Patient Health Questionnaire-9 (PHQ-9) ≥ 8 (Manea et al., 2012). Depression characteristics are presented in Table S4.

Results of these analyses are reported in Table S7. When depression was included in the statistical model, significant negative effects of PTSD on volumes of left lobule VIIB, $b = -119.92$, $t = -3.051$, $p_{\text{-FDR}} = 0.014$, right lobule VIIB, $b = -121.60$, $t = -2.907$, $p_{\text{-FDR}} = 0.021$, and vermal lobule VI, $b = -24.015$, $t = -2.759$, $p_{\text{-FDR}} = 0.025$, were retained. The effect of PTSD on total cerebellum volume did not survive correction for multiple comparisons ($p_{\text{-FDR}} = 0.086$).

Although primary analyses revealed a significant effect of PTSD symptom severity on lobule X volume, there was a significant effect of MDD diagnosis, $b = -8.282$, $t = -2.386$, $p_{\text{-FDR}} = 0.038$, on right lobule X volume, while the effect of PTSD (diagnosis or severity) was non-significant. There was also a significant effect of MDD on corpus medullare volume, $b = -209.30$, $t = -2.513$, $p_{\text{-FDR}} = 0.012$.

**Alcohol use disorder (AUD).** To covary for potential effects of alcohol use disorder (AUD) on total and subregional cerebellar volumes, we performed linear mixed effects models that included fixed effects of age, gender, intracranial volume, PTSD diagnosis, and AUD diagnosis. AUD diagnosis was coded as a binary variable (0=no current alcohol use disorder, 1=current alcohol use disorder). Group status was determined by clinical interview that indicated diagnostic criteria were met for alcohol abuse or dependence (DSM-IV) or alcohol use disorder (DSM-5). Cut-off scores were used to derive group status for subjects without diagnosis but with self-report symptom severity scales from the Alcohol Use Identification Test (AUDIT; Saunders et al., 1993). The AUDIT has a recommended cut-off score of 8 to indicate problematic alcohol use (Allen et al., 1997; Saunders et al., 1993). For these sites, subjects with an AUDIT score ≥8 were coded as "1" and subjects ≤7 were coded as "0." Subjects from sites where alcohol use disorder was a clear exclusion criterion were coded as "0." Sample characteristics for alcohol use disorder are presented in Table S5.

Results of these analyses are reported in Table S8. When AUD was included in the statistical model, there was a significant effect of PTSD diagnosis only on volume of vermal lobule VI, $b = -39.913$, $t = -2.670$, $p_{\text{-FDR}} = 0.035$. There were no significant effects of alcohol use disorder on total and subregional cerebellar volumes.

**Childhood trauma (CTQ).** Additional analyses covaried for potential effects of childhood trauma on total and subregional cerebellar volumes. Cohorts that included administration of the Childhood Trauma Questionnaire (CTQ; Bernstein & Fink, 1998), a retrospective self-report inventory of physical, sexual, and/or emotional abuse during childhood were included in these analyses (n=15 sites). We performed linear mixed effects models that included fixed effects of age, gender, intracranial volume, PTSD diagnosis, and total CTQ severity. Sample CTQ characteristics are reported in Table S5. Subjects with PTSD reported significantly more symptoms of childhood maltreatment than control subjects, $t(1138.7) = -11.26$, $p < .001$.

Results of these analyses are reported in Table S9. When CTQ total severity was included in the statistical model, there were no significant effects of either PTSD diagnosis or childhood trauma on cerebellum volumes after corrections for multiple comparisons. However, PTSD symptom severity was associated with smaller vermis VI ($p_{\text{-FDR}} = 0.022$) and total cerebellum volume ($p_{\text{-FDR}} = 0.036$).

**Table S1:** *Site/study inclusion and exclusion criteria*

| Site | PI(s) | Location(s) | Inclusion criteria | Exclusion criteria |
|---|---|---|---|---|
| **ADNI-DoD** | Thompson | *Marina del Rey, CA, USA* | All: Vietnam War veterans 50-90 years of age, must live within 150 miles of the closest ADNI clinic<br><br>PTSD: Must meet SCID-I (for DSM-IV-TR) criteria for current/chronic PTSD: current CAPS-IV>49; current PTSD symptoms related to a Vietnam War related trauma<br><br>Control: Must be comparable in age, gender, and education with TBI and PTSD groups. May be receiving VA disability payments for something other than TBI or PTSD – or no disability at all. | All: Mild Cognitive Impairment/Dementia; Documented or self-report history of mild/moderate severe TBI; Any history of head trauma associated with persistent cognitive complaints or loss of consciousness >5minutes; History of psychosis, bipolar disorder, alcohol and/or substance abuse/dependence within past 5 years; contraindications to MRI, lumbar puncture, PET scan; unstable medical conditions (e.g., hepatic, renal, pulmonary, metabolic diseases);<br><br>Control: MCI/Dementia; Current or lifetime presence of PTSD (DSM-IV-TR criteria or a CAPS-IV>30) |
| **AMC Amsterdam** | Olff<br>Veltman | *Amsterdam, Netherlands* | All: Police officers 18-65 years of age who are eligible for MRI<br><br>PTSD: current PTSD diagnosis, with CAPS ≥ 45.<br><br>Controls: exposure to at least one traumatic event (according to DSM-IV A1 criterion), with CAPS < 15 | General: History of neurological disorders, any severe or chronic systemic disease or unstable medical condition (including endocrinological disorders), use of psychotropic medications.<br><br>Females: pregnancy or breastfeeding.<br><br>PTSD: current psychotic disorder, substance-related disorder, severe personality disorder, severe major depressive disorder (MDD) (i.e., involving high suicidal risk and/or psychotic symptoms) or current suicidal risk.<br><br>Controls: any current Axis-1 disorder and lifetime history of PTSD or MDD |
| **Beijing** | Li | *Beijing, China* | Individuals 18-65 years of age who personally experienced Wenchuan earthquake in 2008 and are right-handed | Intellectual disability; major psychosis (e.g., schizophrenia and organic mental disorders); drug or alcohol abuse; history of head trauma or surgery; metallic embedded object in body; claustrophobia; exposure to other trauma events from time of the disaster to the time of the study. |
| **Cape Town** | Stein<br>Ipser | *Cape Town, South Africa* | Women 18-65 years of age who speak English, Afrikaans, or Xhosa | Intellectual disability; critical medical condition; current psychotic episode/disorder; contraindications for MRI |
| **Columbia** | Zhu<br>Neria | *New York City, NY, USA* | All: Males or females 18-60 years of age able to give consent, fluent in English<br><br>PTSD: Experience of a traumatic event or events during lifetime; current DSM-V Criterion A for PTSD | All: history of psychosis, bipolar disorder, or dementia; significant depression (HAM-D>25); suicidality; recent substance/alcohol dependence (past 6 months) or abuse (past 2 months); psychotropic medication usage within past 4 weeks (e.g., antipsychotics, antidepressants, mood stabilizers, or stimulants); triptan anti-migraine medications; β-blockers; pregnancy; MRI contraindications; serious and/or unstable/untreated medical illness (e.g., stroke, brain tumor, demyelinating disease)<br><br>Controls: Current or lifetime history of major psychiatric diagnosis, e.g., major depressive disorder, psychotic disorder, bipolar disorder, obsessive compulsive disorder (OCD), PTSD, panic disorder, agoraphobia, eating disorder or alcohol/substance use disorder; history of DSM-5 criterion A1 trauma exposure; HAM-D>7 |

| | | | | |
|---|---|---|---|---|
| **Duke** | Morey | *Durham, NC, USA* | Veterans 18-65 years of age, fluent in English, free of implanted metal objects or metal shards in eyes | Axis I psychiatric conditions other than PTSD or MDD; current substance abuse or lifetime substance dependence (other than nicotine); high risk for suicide, claustrophobial; neurological disorders; learning disability or developmental delay; major medical conditions |
| **Emory GTP** | Stevens<br>Fani | *Atlanta, GA, USA* | Individuals 18-65 years of age who speak English and have endorsed at least 1 criterion A trauma | Current psychotic symptoms or bipolar disorder; current substance or alcohol dependence; history of head trauma; psychoactive medication usage; current illegal drug use (verified with urine drug screen within 24 hours of scan) |
| **Ghent** | Mueller | *Ghent, Belgium* | All: Individuals 18-60 years of age who speak Dutch and have normal or corrected-to-normal vision.<br><br>Childhood abuse: endorsed physical, sexual, and/or emotional abuse prior to age 17.<br><br>No abuse: no history of childhood trauma, no lifetime history of abuse-related trauma. | All: contraindications to MRI.<br><br>No abuse: lifetime history of abuse-related trauma. |
| **Groningen** | Daniels | *Groningen, Netherlands* | Civilian women 20-60 years of age with a current PTSD diagnosis, sufficient proficiency in German, MRI compatible | Neurologic disorders; history of substance abuse or dependence (past 6 months); history of head injury; cerebral incidental findings verified by a neuroradiologist after the MR scan; usage of benzodiazepines, tricyclic antidepressants, or anticonvulsants; primary borderline personality disorder; other current diagnosis of Axis I disorder |
| **LIMBIC-CENC** | Dennis<br>Tate<br>Cifu<br>Walker<br>Wilde | *Richmond, VA*<br>*Houston, TX*<br>*Tampa, FL*<br>*San Antonio, TX*<br>*Ft. Belvoir, FL*<br>*Portland, OR*<br>*Minneapolis, MN*<br>*USA* | Veterans with history of deployment in Operation Enduring Freedom (OEF), Operation Iraqi Freedom (OIF), Operation New Dawn (OND), or follow-up conflicts; history of combat exposure (score>1 on any item in Deployment Risk and Resiliency Inventory Section D [DRRI-2-D]) | History of moderate to severe TBI; history of major neurologic disorder with significant decrease in functional status and/or loss of ability for independent living; severe psychiatric disorder (e.g., schizophrenia) |
| **Mannheim** | Schmahl<br>Herzog | *Mannheim, Germany* | PTSD: Women aged 18-65 years with PTSD related to index trauma of sexual or physical abuse before the age of 18 years; 3+ criteria of Borderline Personality Disorder (including affective instability); able to attend weekly therapy sessions for one year<br><br>Trauma-exposed control: Women aged 18-65 years; Childhood sexual or physical abuse before the age of 18 years.<br><br>Healthy control: Women aged 18-65 years | PTSD: planned absence >4 weeks from therapy |
| **Masaryk** | Rektor<br>Riha | *Brno, Czech Republic* | Holocaust survivors and subsequent generations exposed to varying levels of trauma who are 15-65 years of age. Must be fluent in Czech/Slovak, able to provide informed consent, and have an MMSE score >26. | History of neurological or psychotic disorder; contraindication to MRI |
| **McLean 1** | Kaufman<br>Ressler | *Boston, MA, USA* | Women 18-60 years of age with a history of childhood maltreatment who speak English; must have legal and mental competency, Normal or Corrected Vision. | Delirium secondary to medical illness; History of neurological conditions that may cause significant psychiatric symptomatology (e.g., dementia); Any contraindication to MR scans, including claustrophobia, pregnancy, metal implants, etc.; Current alcohol or substance use disorder (within the last month); A history of schizophrenia or other psychotic disorder; History of head injury or loss of consciousness for longer than 5 min (including concussion); pregnancy |

| | | | | |
|---|---|---|---|---|
| **McLean 2** | Rosso | *Boston, MA, USA* | Civilians aged 20-50 years old; right-handed; DSM-IV diagnosis consistent with group assignment; ability to provide written informed consent | Medical condition that would confound results; history of seizures or head trauma with loss of consciousness; exposure to psychotropic medications within 4 weeks of study (8 weeks for fluoxetine); contraindications to MRI; positive urine toxicology or HCG status on scan day; history of psychotic disorder, bipolar disorder, eating disorder, intellectual disability, or pervasive developmental disorder; lifetime history of DSM-IV non-PTSD anxiety disorder |
| **Michigan** | Liberzon | *Ann Arbor, MI, USA* | All: Male combat veterans and civilians who are 18-55 years of age and eligible for MRI.<br><br>PTSD: current PTSD diagnosis with CAPS-IV ≥50. Trauma-exposed controls: exposure to ≥1 traumatic event (per DSM-IV A1 criterion) with CAPS-IV <15. Community controls: CAPS-IV < 15. | History of neurological disorders, any severe or chronic disorder, current alcohol or drug abuse and/or dependence. |
| **Milwaukee** | Larson | *Milwaukee, WI, USA* | Civilians aged 18-60 years; exposure to DSM-5 A1 criterion trauma; high risk for PTSD (score ≥3 OR item 2 rated ≥3 on Predicting PTSD Questionnaire, Rothbaum et al., 2014); English speaking; ability to schedule baseline study visit within 30 days of traumatic injury | Glasgow Coma Scale score ≤ 13 (i.e., moderate to severe traumatic brain injury); on police hold; contraindication to MRI; pregnancy (or planned pregnancy within 6 months); intentional self-inflicted injury; severe vision or hearing impairment; history of psychotic or manic symptoms, or neurologic condition (e.g., seizures, spinal cord injury); currently on antipsychotic medication; clear evidence of substance use disorder |
| **Minnesota** | Lissek | *Minneapolis, MN, USA* | Individuals 18-65 years of age with history of combat-related trauma | Current of past history of psychosis, bipolar disorder, delirium, dementia, amnestic disorder, or intellectual disability; suicidality; substance use disorder within past six months; pregnancy; current or past medical illnesses that may confound study results or place participant at risk; current use of any medication that alters central nervous system function including antidepressants, benzodiazepines, anti-psychotics, mood-stabilizers, anti-parkinsonian agents, anti-convulsants, sleep medications, pain medications, and anti-hypertensives; MRI contraindications |
| **Missouri** | Bruce | *St. Louis, MO, USA* | Patients: women with current DSM-IV diagnosis of PTSD related to interpersonal trauma, with CAPS-IV ≥45.<br><br>Controls: no history of DSM-IV criterion A trauma exposure | All: history of psychosis or bipolar disorder; current psychotropic medication usage; current alcohol or substance use disorder; history of significant head trauma or neurologic condition; contraindication to MRI.<br><br>Controls: current mood or anxiety disorder. |
| **Münster** | Straube Hofmann | *Münster, Germany* | Individuals who have experienced IPV trauma; right-handed; normal or corrected-to-normal vision | |
| **Nanjing** | Qi | *Nanjing, China* | Individuals 40-70 years of age who have experienced the death of their only child within past 10 years | Current psychiatric disorder (except PTSD, MDD, GAD); history of brain injury or other serious medical or neurological condition; MRI contraindications; left-handedness |
| **South Dakota** | Baugh Fercho | *Vermillion, SD, USA* | OIF/ OEF/OND veterans | Current or previous seizure history; current crisis-related issues such as serious self-injurious behavior, psychosis, or substance dependence (excluding alcohol dependence); report of traumatic brain injury using the Traumatic Brain Injury Checklist (Hoge et al., 2008); contraindications to fMRI (metal objects in body, claustrophobia) |

| | | | | |
|---|---|---|---|---|
| | | | OEF/OIF Veterans 18 and 65 years of age; fluent in English; able to provide informed consent. | History of psychotic, bipolar, or substance dependence (within 3 months for PTSD group or lifetime for controls), history of neurological disorder, moderate to severe traumatic brain injury, claustrophobia, regular use of benzodiazepines, opiates, thyroid medications, or other CNS medications.<br><br>Trauma-exposed healthy controls: history of any Axis I psychiatric disorder, including PTSD. |
| **Stanford** | Etkin<br>Maron-Katz | *Palo Alto, CA, USA* | All: Community-dwelling adults 18-60 years of age who are fluent in English<br><br>Patients: Individuals experiencing chronic (>3 months) moderate to severe anxiety or depression (indicated by a score >10 on the PHQ-9 [excluding suicide item] OR a score >10 on the GAD-7 scale) - must express interest in seeking treatment for psychiatric symptoms.<br><br>Controls: PHQ-9 and GAD ≤ 4. | All: contraindications to MRI or TMS; current participation in psychiatric treatment; history of neurological disorder, brain surgery, electroconvulsive or radiation treatment, brain hemorrhage or tumor, stroke, epilepsy, hypo- or hyper-thyroidism; medication use that substantially reduces seizure threshold to TMS (e.g., olanzapine, chlorpromazine, lithium) and unwillingness or inability to safely withdraw at least two weeks prior to TMS appointment; medications that interfere with blood flow (e.g., opiates, antihypertensive medication); insufficiently controlled thyroid dysfunction; current substance dependence (within past 3 months); refusal to abstain from illicit drug use for study duration; refusal to abstain from alcohol within 24 hours of MRI scans; pregnancy; prior exposure to deep brain stimulation, rTMS, or tDCS therapies; significant traumatic brain injury (indicated by loss of consciousness, post-trauma amnesia, imaging findings, penetrating brain injury); history of psychotic or manic symptoms. |
| **Toledo** | Wang | *Toledo, OH, USA* | Motor vehicle accident (MVA) survivors transported to the University of Toledo Emergency department, or to a ProMedica emergency medicine department. | Pregnancy; under the influence of alcohol or drugs at the time of MVA; major injuries, moderate to severe traumatic brain injury; major medical illnesses; contraindication to MRI |
| | | | Ohio National Guard and Reserve soldiers 18-50 years of age who were deployed in OEF or OIF – must have met Ohio National Guard Study characteristics and able to provide informed consent. | History of psychosis, bipolar disorder, or neurologic condition; current substance dependence; intellectual disability or developmental disorder; contraindication to MRI; current use of antipsychotic medication. |
| **Tours** | Quide<br>El Hage | *Tours, France* | Female survivors of sexual assault (within past month)<br><br>Control: no history of interpersonal violence exposure | History of head injury, substance abuse, current use of psychotropic medication, medical conditions with confounding effects on brain function (e.g., epilepsy, brain tumor), contraindication to MRI |
| **VA Minneapolis** | Disner<br>Davenport | *Minneapolis, MN, USA* | OEF and/or OIF veteran 22-60 years of age who had been exposed to combat during their deployment(s). | Current psychosis; current DSM-IV substance abuse or dependence other than alcohol, caffeine, or nicotine; moderate or severe traumatic brain injury; neurologic condition other than TBI; current unstable medical condition that would likely affect brain function (e.g., uncontrolled diabetes); significant imminent risk of suicidal or homicidal behavior. |
| **VA Waco** | May<br>Gordon | *Waco, TX, USA* | Veterans 18-60 years of age | Seizure disorder, dementia; MRI contraindications |
| | | | Veterans 18-60 years of age with a clinical diagnosis of TBI in their VA medical record | Diagnosis of schizophrenia, schizoaffective disorder, bipolar disorder type I, severe substance use disorder, or a high risk of suicide; absence of qEEG more than 2SDs outside of population means of healthy age-matched controls; MRI contraindications |
| **VA West Haven** | Abdallah | *West Haven, CT, USA* | Combat-exposed US Veterans aged 21-65 both with and without PTSD, fluent in English | History of psychotic disorder, bipolar depression, or neurologic/neurodevelopmental disorder (including learning disorders, attention- |

| | | | | |
|---|---|---|---|---|
| | | | | deficit hyperactivity disorder, moderate to severe traumatic brain injury, epilepsy, brain tumor); contraindication to MRI |
| **Vanderbilt** | Blackford | *Nashville, TN, USA* | OEF/OIF/OND Veterans 18-50 years of age who are fluent in English | Psychoactive medication usage in past 6 weeks; participated in psychotherapy within the past month; current substance use disorder (>6 month remission); positive urine drug or alcohol breath screen on MRI study day; history of psychotic or bipolar disorder, traumatic brain injury, or significant medical (e.g., cancer, HIV) or neurological illness (e.g., stroke, brain tumor, multiple sclerosis, epilepsy); contraindication to MRI<br><br>Trauma-exposed controls: lifetime diagnosis of PTSD; symptoms of hypervigilance<br><br>Healthy controls: any trauma exposure |
| **VETSA** | Franz | *San Diego, CA, USA* | Brothers 50-59 years of age from the Vietnam Era Twin Registry (VETR); both in the US military at some point between 1965 and 1975; both brothers willing to participate<br><br>*1 brother from each pair was randomly selected for the present study analyses | MRI contraindications |
| **Wisconsin 1** | Cisler | *Madison, WI, USA* | Women aged 21-50 with and without history of interpersonal violence – must be fluent in English | History of psychosis, medication changes within past 4 weeks, cognitive impairment, current substance or alcohol use disorder |
| **Wisconsin 2** | Grupe | *Madison, WI, USA* | Adults 18-50 with exposure to 1+ life-threatening war zone trauma events; capable of giving informed consent and fluent in English; clear evidence of war zone trauma exposure in Iraq or Afghanistan since 2001 (e.g., Combat Action Ribbon [Marines], Combat Infantry Badge [Army]); stable pharmacological or psychotherapeutic treatment for at least 8 weeks prior to beginning of study | Weight >352 pounds or over; pregnancy or current breastfeeding; Metallic implants such as prostheses or aneurysm clip, or electronic implants such as cardiac pacemakers; Neurological or serious medical condition; History of seizures or seizure disorder; Moderate or severe traumatic brain injury; Current active substance dependence or dependence within 3 months (other than nicotine); bipolar disorder, schizophrenia, schizoaffective disorder, psychotic disorder NOS, delirium, or any DSM-IV cognitive disorder; Severe psychiatric instability or severe situational life crises, (e.g., suicidality, homicidality); extensive experience in yoga or meditation; Current use of benzodiazepines and beta-blockers |
| **Yale** | Harpaz-Rotem | *New Haven, CT, USA* | Individuals 21-60 years of age; at least one deployment on combat tour | Diagnosis of bipolar disorder or psychotic disorder; current benzodiazepine use; a history of ADHD, learning disorder, moderate or severe traumatic brain injury (TBI), brain tumor, epilepsy, or a neurological disorder; current inpatient status; MRI contraindication. |

**Table S2:** *Scan parameters by site*

| Site | Scanner | Field strength | # head coil channels | Sequence | Voxel size (mm) | FOV (mm) | Orientation | Repetition time (TR) | Echo time (TE) | Flip angle |
|---|---|---|---|---|---|---|---|---|---|---|
| **ADNI DoD** | GE Discovery MR750w | 3T | 40 | FSPGR | 1x1x1.2 | 256x256 | Sagittal | 7652 | 3.10 | 11 |
| | GE Discovery MR750 | 3T | 8 | SPGR | 1x1x1.2 | 256x256 | Sagittal | 6984 | 2.85 | 11 |
| | GE Signa HDxt | 3T | 8 | SPGR | 1x1x1.2 | 256x256 | Sagittal | 7340 | 3.04 | 11 |
| | Siemens TIM Trio | 3T | 12 | MPRAGE | 1x1x1.2 | 256x256 | Sagittal | 2300 | 2.98 | 9 |
| **AMC Amsterdam** | Philips Achieva | 3T | 32 | FAST MPRAGE | 1x1x1 | 240x188 | Axial | 8200 | 3.8 | 8 |
| **Beijing** | Philips Achieva | 3T | 8 | EPI | 1x1x1 | 220x220 | Axial | 8500 | 3.7 | 8 |
| **Cape Town** | Siemens Skyra | 3T | 4 | MPRAGE | 1x1x1.5 | 256x256 | Sagittal | 2530 | 1.69 / 3.55 / 5.41 / 7.27 | 7 |
| | Siemens Allegra | 3T | 4 | MPRAGE | 1x1x1.5 | 256x256 | Sagittal | 2000 | 1.53 / 3.21 / 4.89 / 6.57 | 20 |
| **Columbia** | GE Signa Excite | 1.5T | 8 | SPGR | 3.5x3.5x2.2 | 224x224 | Axial | 3000 | 3.0 | 84 |
| **Duke** | GE Discovery MR750 | 3T | 8 | FSPGR BRAVO | 1x1x1 / 0.9375x0.9375x1 | 256x256 / 240x240 | Axial | 8160 | 3.2 | 12 |
| | GE Signa Excite | 3T | 8 | FSPGR BRAVO | 0.9375x0.9375x1 | 240x240 | Axial | 8148 / 7840 / 8160 | 3.22 / 2.9 | 12 |
| | Philips Ingenia | 3T | 8 | 3D TFE SENSE | 0.9375x0.9375x1 | 240x240 | Axial | 8.148 | 3.73 | 8 |
| **Emory GTP** | Siemens TIM Trio | 3T | 12 | MPRAGE | 1x1x1 | 224x256 | Axial | 2600 | 3.02 | 8 |
| **Ghent** | Siemens TIM Trio | 3T | 32 | MPRAGE | 1x1x1 | 256x256 | Transversal | 2250 | 4.18 | 9 |
| **Groningen** | Siemens TIM Trio | 3T | 12 | MPRAGE | 1x1x1 | 256x256 | Sagittal | 1900 | 2.52 | 9 |
| **LIMBIC-CENC** | Philips Ingenia | 3T | na | MPRAGE | 1x1x1.2 | 256x256 | Sagittal | 6.78 | 3.16 | 9 |
| | Siemens TIM Trio | 3T | na | MPRAGE | 1x1x1.2 | 240x256 | Sagittal | 2300 | 2.96 | 9 |
| | GE Signa HDxt | 3T | na | SPGR | 1x1x1.2 | 256x256 | Sagittal | 6.28 | 2.78 | 11 |
| | Siemens Verio/Skyra Fit | 3T | na | MPRAGE | 1x1x1.2 | 240x256 | Sagittal | 2300 | 2.98 | 9 |
| | GE Discovery MR750 | 3T | na | SPGR | 1x1x1.2 | 256x256 | Sagittal | 8.156 | 3.18 | 11 |
| | Philips Achieva | 3T | na | MPRAGE | 1x1x1x1.2 | 256x256 | Sagittal | 6.76 | 3.15 | 9 |
| | Siemens Prisma | 3T | na | MPRAGE | 0.8x0.8x0.8 | 300x320 | Sagittal | 2400 | 2.24 | 8 |
| | Siemens Prisma | 3T | na | MPRAGE | 0.8x0.8x0.8 | 300x320 | Sagittal | 2400 | 2.24 | 8 |
| **Mannheim** | Siemens TIM Trio | 3T | 32 | SPGR | 1x1x1 | 192x192 | Axial | 2000 | 3.0 | 80 |
| **Masaryk** | Siemens Prisma | 3T | 64 | MPRAGE | 1x1x1 | 224x224 | na | 2300 | 2.34 | 8 |
| **McLean 1** | Siemens TIM Trio | 3T | 32 | MPRAGE | 1.3x1.3x1.3 | 256x128 | Sagittal | 2530 | 3.31 | 7 |
| **McLean 2** | Siemens TIM Trio | 3T | 12 | MEMPRAGE | 1x1x1 | 256x256 | Sagittal | 2530 | 1.64 / 3.5 / | 10 |

| | | | | | | | | | | |
|---|---|---|---|---|---|---|---|---|---|---|
| | | | | | | | | | 5.36 / 7.22 | |
| **Michigan** | GE Signa Excite | 3T | 8 | IF-FSPGR | 1x1x1 | 256x256 | Axial | 12300 | 5.3 | 9 |
| **Milwaukee** | GE Discovery MR750 | 3T | 32 | SPGR | 1x0.9375x0.9375 | 240x240 | Sagittal | 9800 | 4.6 | 8 |
| **Minnesota** | Siemens Prisma | 3T | 32 | na | 0.9x0.9x0.9 | na | na | na | na | na |
| **Missouri** | Siemens TIM Trio | 3T | 12 | MPRAGE | 1x1x1 | 256x256 | Sagittal | 2400 | 3.13 | 8 |
| **Münster** | Siemens Prisma | 3T | 32 | MPRAGE | 1x1x1 | 256x256 | Sagittal | 2130 | 2.28 | 8 |
| **Nanjing** | GE Discovery MR750 | 3T | 8 | FSPGR BRAVO | 1x1x1 | 240x240 | Axial | 8208 | 3.22 | 12 |
| **South Dakota** | Siemens Skyra | 3T | 32 | MPRAGE | 1x1x1 | 240x240 / 256x256 | Sagittal | 1900 | 2.13 | 9 |
| **Stanford** | GE Discovery MR750 | 3T | 8 | SPGR | 1.5x0.9x1.1 | 220x220 / 240x240 | Coronal | 8000 / 8600 | 3.6 / 3.4 | 15 |
| **Toledo** | GE SignaX | 3T | 8 | SPGR | 1x1x1 | 256x256 | Axial | 8200 | 3.2 | 12 |
| **Tours** | Siemens Verio | 3T | 12 | na | 1x1x1 | 256x256 | Sagittal | 1900 | 2.48 | 9 |
| **VA Minneapolis** | Siemens Tim Trio | 3T | 12 | MPRAGE | 1x1x1 | 256x256 | Coronal | 2530 | 3.7 | 7 |
| **VA Waco** | Philips Achieva | 3T | 16 | MPRAGE | 0.9x0.9x0.9 | 256x256 | Sagittal | 7256 | 2.77 | 12 |
| **VA West Haven** | Siemens TIM Trio | 3T | 32 | MPRAGE | 1x1x1 | 256x256 | Sagittal | 2530 | 2.71 | 7 |
| **Vanderbilt** | Philips Intera | 3T | 32 | na | 0.8x0.8x0.9 | 256x256 | Sagittal | 9000 | 4.6 | 9 |
| **VETSA** | Siemens TIM Trio | 3T | 32 | MPRAGE | 1x1x1.2 | 256x256 | Sagittal | 2170 | 4.3 | 7 |
| | GE Discovery MR750 | 3T | 8 | FSPGR | 1x1x1.2 | 240x240 | Sagittal | 8084 | 3.16 | 8 |
| **Wisconsin 1** | Philips Achieva X-Series | 3T | 32 | MPRAGE | 1x1x1 | 256x256 | Sagittal | 7500 | 3.7 | 9 |
| | GE Discovery MR750 | 3T | 8 | MPRAGE | 1x1x1 | 256x256 | Axial | 8200 | 3.2 | 12 |
| **Wisconsin 2** | GE Discovery X750 | 3T | 8 | MPRAGE | 1x1x1 | 256x256 | Sagittal | 1900 | 2.5 | 9 |
| **Yale** | Siemens TIM Trio | 3T | 12 | MPRAGE | 1x1x1 | 256x256 | Sagittal | 2500 | 2.77 | 7 |

*Notes:* BRAVO, inversion preparation of a fast low-angle spoiled gradient recall; EPI, echo planar image; FAST, fast low-angle shot; FOV, field-of-view; GE, General Electric; MEMPRAGE, multi-echo magnetization-prepared rapid gradient echo; MPRAGE, magnetization-prepared rapid gradient echo; na, not available; SPGR, spoiled gradient recall; TFE SENSE, turbo field echo sensitivity encoding; TR, repetition time; TE, echo time

**Table S3:** *Cerebellum parcellation quality control ratings by site.*

| Site | Mean | Standard Deviation |
|---|---|---|
| ADNI DoD | 1.862 | 0.944 |
| Amsterdam AMC | 1.027 | 0.232 |
| Beijing | 1.034 | 0.237 |
| Cape Town | 1.155 | 0.453 |
| Columbia | 1.349 | 0.656 |
| Duke | 1.228 | 0.507 |
| Emory GTP | 1.103 | 0.411 |
| Ghent | 1.090 | 0.379 |
| Groningen | 1.512 | 0.675 |
| LIMBIC-CENC | --- | --- |
| Mannheim | 1.673 | 0.774 |
| Masaryk | 1.132 | 0.432 |
| McLean 1 | 1.025 | 0.225 |
| McLean 2 | 1.054 | 0.324 |
| Michigan | 1.032 | 0.252 |
| Milwaukee | 1.571 | 0.765 |
| Minnesota | 1.032 | 0.178 |
| Missouri | 1.121 | 0.412 |
| Münster | 1.298 | 0.623 |
| Nanjing | 1.168 | 0.544 |
| South Dakota | 1.390 | 0.622 |
| Stanford | 1.623 | 0.878 |
| Toledo | 1.063 | 0.334 |
| Tours | 1.333 | 0.612 |
| VA Minneapolis | 1.037 | 0.228 |
| VA Waco | 1.450 | 0.642 |
| VA West Haven | 1.354 | 0.738 |
| Vanderbilt | 1.360 | 0.631 |
| VETSA | 1.146 | 0.454 |
| Wisconsin 1 | 1.029 | 0.217 |
| Wisconsin 2 | 2.000 | 0.918 |
| Yale | 1.082 | 0.363 |
| | **1.269** | **0.224** |

*Note.* Subjects' cerebellar parcellation outputs were scored as 1 "good," 2 "acceptable," or 3 "failed/poor" based on visual examination. Fleiss' kappa showed moderate inter-rater agreement on segmentation quality, $\kappa = .469$ (95% CI, .444-.494), $p < .001$.Cerebellar parcellation for LIMBIC-CENC sites (n=1045) was performed at the University of Utah. Ratings are unavailable for these sites due to differences in quality control procedures; however, consistent with procedures at Duke University, quality control of these scans also included quantitative outlier removal and visual examination of outputs for segmentation errors or failure.

**Table S4:** *Depression diagnosis and severity by site.*

| | Depression Diagnosis | | Diagnostic Tool | Depression Severity | Severity Tool |
|---|---|---|---|---|---|
| | *MDD* | *Ctrl* | | *M % (SD)* | |
| ADNI | 2 | 99 | GDS | 16.76 (18.34) | GDS |
| Amsterdam | 8 | 65 | HADS | 27.16 (28.70) | HADS-D |
| Beijing | 51 | 36 | CES-D | 33.82 (17.31) | CES-D |
| Cape Town | 30 | 72 | BDI-II | 22.28 (16.84) | BDI-II |
| LIMBIC-CENC | 667 | 372 | PHQ-9 | 29.83 (22.81) | PHQ-9 |
| Columbia | 32 | 119 | SCID, HAM-D | 17.33 (16.13) | HAM-D |
| Duke | 71 | 305 | BDI-II | 18.84 (20.19) | BDI-II |
| Emory | 18 | 39 | BDI-II | 24.09 (17.69) | BDI-II |
| Ghent | 11 | 54 | MINI | 17.58 (15.90) | BDI-II |
| Groningen | 3 | 34 | BDI-II | 36.51 (21.42) | BDI-II |
| Mannheim | 37 | 3 | BDI-II | 58.69 (20.91) | BDI-II |
| Masaryk | 49 | 220 | GDS | --- | --- |
| McLean 1 | 44 | 34 | BDI-II | 31.40 (24.69) | BDI-II |
| McLean 2 | 15 | 79 | BDI-II | 9.59 (13.83) | BDI-II |
| Michigan | 25 | 15 | MINI | 36.80 (30.51) | DASS-21 |
| Minnesota VA | 86 | 155 | SCID | --- | --- |
| Münster | 10 | 33 | BDI-II | 15.95 (18.92) | BDI-II |
| Nanjing | 15 | 117 | SCID | --- | --- |
| South Dakota | 28 | 86 | CES-D | 19.97 (17.08) | CES-D |
| Stanford | --- | --- | --- | --- | --- |
| Toledo | 24 | 48 | CES-D, DASS-21 | 29.54 (26.25) | CES-D, DASS-21 |
| Tours | 3 | 36 | BDI-SF | 11.11 (15.28) | BDI-SF |
| Minnesota | 8 | 54 | SCID | 16.99 (15.25) | BDI-II |
| Missouri | 43 | 21 | BDI-II | 38.24 (17.04) | BDI-II |
| Wisconsin 1 | 21 | 83 | SCID | 30.44 (21.96) | BDI-II |
| Wisconsin 2 | 12 | 12 | BDI-II | 40.15 (17.85) | BDI-II |
| Milwaukee | 11 | 59 | MINI | 22.38 (26.95) | DASS-21 |
| Vanderbilt | 4 | 42 | MINI | 9.11 (12.32) | BDI-II |
| VETSA | 20 | 170 | CES-D | 10.53 (11.94) | CES-D |
| Waco VA | 42 | 23 | BDI-II | 32.41 (20.78) | BDI-II |
| West Haven VA | 14 | 41 | SCID, BDI-II | 31.26 (19.20) | BDI-II |
| Yale | 17 | 54 | BDI-II | 16.19 (18.95) | BDI-II |
| **Overall** | **1421** | **2580** | **---** | **24.71 (22.17)** | **---** |

*Notes:* BDI-II, Beck Depression Inventory-II; BDI-SF, Beck Depression Inventory-Short Form; CES-D, Center for Epidemiological Studies Depression Scale; DASS-21, Depression Anxiety Stress Scales-21 item; GDS, Geriatric Depression Scale; HADS, Hospital Anxiety and Depression Scale; HAM-D, Hamilton Depression Rating Scale; MINI, Mini International Neuropsychiatric Interview; PHQ-9, Patient Health Questionnaire-9; SCID, Structured Clinical Interview for DSM

**Table S5:** *Alcohol use and childhood trauma characteristics by site.*

| | Alcohol Use | | Diagnostic Tool | CTQ Total |
|---|---|---|---|---|
| | *AUD* | *Ctrl* | | *M (SD)* |
| ADNI-DoD | 37 | 52 | Interview | --- |
| Amsterdam AMC | --- | --- | --- | --- |
| Beijing | --- | --- | --- | --- |
| Cape Town | 26 | 78 | ASSIST | 39.37 (14.12) |
| LIMBIC-CENC | --- | --- | --- | --- |
| Columbia | --- | --- | --- | 43.38 (19.25) |
| Duke | 16 | 100 | AUDIT | 61.68 (16.40) |
| Emory | --- | --- | --- | 42.92 (17.15) |
| Ghent | 2 | 63 | MINI | --- |
| Groningen | 5 | 32 | SCID | 85.09 (12.24) |
| Mannheim | 0 | 40 | Exclusion | 79.43 (19.03) |
| Masaryk | --- | --- | --- | --- |
| McLean 1 | 11 | 66 | SCID | 61.19 (29.20) |
| McLean 2 | --- | --- | --- | 50.22 (22.10) |
| Michigan | 0 | 62 | Exclusion | --- |
| Minnesota VA | --- | --- | --- | --- |
| Münster | --- | --- | --- | --- |
| Nanjing | 0 | 132 | Exclusion | --- |
| South Dakota | --- | --- | --- | --- |
| Stanford | --- | --- | --- | 53.62 (25.19) |
| Toledo | 8 | 66 | MINI | 51.76 (16.48) |
| Tours | 0 | 45 | MINI | --- |
| Minnesota | 0 | 62 | SCID | --- |
| Missouri | --- | --- | --- | --- |
| Wisconsin 1 | 0 | 104 | Exclusion | 55.91 (26.43) |
| Wisconsin 2 | --- | --- | --- | --- |
| Milwaukee | --- | --- | --- | 44.67 (15.46) |
| Vanderbilt | 0 | 46 | Exclusion | 40.91 (12.76) |
| VETSA | 174 | 16 | Interview | --- |
| Waco VA | --- | --- | --- | 51.13 (21.38) |
| West Haven VA | --- | --- | --- | --- |
| Yale | --- | --- | --- | 47.56 (17.25) |
| **Overall** | **279** | **964** | **---** | **52.91 (22.69) n=1190** |

*Notes:* ASSIST, Alcohol Smoking and Substance Involvement Screening; AUDIT, Alcohol Use Disorders Identification Test; AUD, Alcohol Use Disorder; CTRL, Control; CTQ, Childhood Trauma Questionnaire; MINI, Mini International Neuropsychiatric Interview; SCID, Structured Clinical Interview for DSM

**Table S6:** *Race and ethnicity by site.*

| | Race (%) | | | | | Ethnicity (%) | | |
|---|---|---|---|---|---|---|---|---|
| **Site** | *White* | *Black* | *Asian* | *Mixed Race* | *Other or Not Reported* | *Hispanic* | *Non-Hispanic* | *Not Reported* |
| ADNI DoD | 83.5 | 7.8 | 2.9 | 1.9 | 3.9 | 7 | 41 | 55 |
| Amsterdam AMC | 100 | --- | --- | --- | --- | 2 | 71 | --- |
| Beijing | --- | --- | 100 | --- | --- | --- | 87 | --- |
| Cape Town | --- | 40.6 | --- | 59.4 | --- | --- | --- | 100 |
| Columbia | 29.8 | 38.4 | 4.0 | --- | 27.8 | 44 | 107 | --- |
| Duke | 47.1 | 38.6 | --- | --- | 14.4 | 17 | 265 | 94 |
| Emory GTP | --- | 100 | --- | --- | --- | --- | --- | 100 |
| Ghent | 100 | --- | --- | --- | --- | --- | --- | 100 |
| Groningen | 100 | --- | --- | --- | --- | 2.7 | 97.3 | --- |
| LIMBIC-CENC | 73.7 | 17.5 | 1.7 | --- | 7.1 | 17.7 | 80.7 | 1.6 |
| Mannheim | 100 | --- | --- | --- | --- | --- | 100 | --- |
| Masaryk | 100 | --- | --- | --- | --- | --- | --- | 100 |
| McLean 1 | 87 | 5.2 | 6.5 | --- | 1.3 | 6.5 | 93.5 | --- |
| McLean 2 | 68.1 | 13.8 | 13.8 | --- | 4.3 | 10.6 | 89.4 | --- |
| Michigan | 87.1 | 6.5 | 1.6 | 3.2 | 1.6 | --- | 100 | --- |
| Milwaukee | 31.4 | 48.6 | 1.4 | 10 | 8.6 | 12.9 | 85.7 | 1.4 |
| Minnesota | 95.2 | --- | 1.6 | 1.6 | 1.6 | 1.6 | 96.8 | 1.6 |
| Missouri | 54.8 | 38.7 | --- | 3.2 | 3.2 | --- | --- | 100 |
| Münster | 100 | --- | --- | --- | --- | --- | --- | 100 |
| Nanjing | --- | --- | 100 | --- | --- | --- | 100 | --- |
| South Dakota | 92.1 | 2.6 | 0.9 | 1.8 | 2.6 | --- | --- | 100 |
| Stanford | 33.8 | 0.7 | 11.7 | 5.5 | 48.3 | 11 | 46.9 | 42.1 |
| Toledo | 53.2 | 19.5 | --- | --- | 27.3 | 1.3 | 19.5 | 79.2 |
| Tours | --- | --- | --- | --- | 100 | --- | --- | 100 |
| VA Minneapolis | 89.2 | 1.7 | 4.1 | 0.4 | 4.6 | --- | --- | 100 |
| VA Waco | 51.1 | 18.5 | 1.1 | 25 | 4.3 | 22.8 | 75 | 2.2 |
| VA West Haven | --- | --- | --- | --- | 100 | --- | --- | 100 |
| Vanderbilt | 78.3 | 10.9 | --- | 10.9 | --- | 2.2 | 97.8 | --- |
| VETSA | --- | --- | --- | --- | 100 | --- | --- | 100 |
| Wisconsin 1 | 90.4 | 16 | 1.1 | --- | 3.2 | 2 | 0 | 102 |
| Wisconsin 2 | 95.8 | --- | --- | --- | 4.2 | --- | 100 | --- |
| Yale | 76.8 | 7.2 | 1.4 | --- | 14.5 | 8.7 | 91.3 | --- |
| **n** | **2559** | **640** | **298** | **116** | **602** | **328** | **2244** | **1643** |
| **% of overall sample** | **60.7%** | **15.2%** | **7.1%** | **2.8%** | **14.3%** | **7.8%** | **53.2%** | **39.0%** |

Due to differences in reported racial categories by site and small sample sizes, categories for which no site reported more than 5 participants were combined with “Other/Not reported.”

**Table S7:** *Effects of PTSD & MDD diagnosis on cerebellar volumes.*

| | | PTSD | | | | | MDD | | | | |
|---|---|---|---|---|---|---|---|---|---|---|---|
| **ROI** | **N** | ***b*** | ***SE*** | ***t*** | ***p*-FDR** | ***d*** | ***b*** | ***SE*** | ***t*** | ***p*-FDR** | ***d*** |
| **Anterior** | | | | | | | | | | | |
| Left I-III | 3971 | -11.459 | 7.283 | -1.573 | 0.348 | -0.050 | 4.438 | 7.753 | 0.572 | 0.999 | 0.018 |
| Left IV | 3952 | -7.063 | 19.993 | -0.353 | 0.724 | -0.011 | -17.366 | 21.284 | -0.816 | 0.623 | -0.026 |
| Left V | 3909 | 8.079 | 17.830 | 0.453 | 0.975 | 0.015 | 0.770 | 18.981 | 0.041 | 0.968 | 0.001 |
| Right I-III | 3972 | -8.280 | 7.690 | -1.077 | 0.423 | -0.034 | 2.510 | 8.174 | 0.307 | 0.999 | 0.010 |
| Right IV | 3949 | -1.410 | 20.930 | -0.067 | 0.946 | -0.002 | 1.366 | 22.278 | 0.061 | 0.999 | 0.002 |
| Right V | 3903 | -41.97 | 19.53 | -2.149 | 0.096+ | -0.071 | -19.95 | 20.74 | -0.962 | 0.999 | -0.032 |
| **Posterior** | | | | | | | | | | | |
| Left Crus I | 3766 | 0.684 | 71.216 | 0.010 | 0.992 | 0.003 | 23.246 | 75.761 | 0.307 | 0.999 | 0.010 |
| Left Crus II | 3899 | -97.57 | 46.95 | -2.078 | 0.133 | -0.067 | -23.81 | 50.10 | -0.475 | 0.999 | -0.015 |
| Left VI | 3952 | -29.26 | 47.67 | -0.614 | 0.943 | -0.020 | 66.38 | 50.65 | 1.311 | 0.999 | 0.042 |
| **Left VIIB** | 3885 | **-119.92** | **39.31** | **-3.051** | **0.014*** | **-0.098** | 23.14 | 41.78 | 0.554 | 0.999 | 0.018 |
| Left VIIIA | 3825 | 4.649 | 39.667 | 0.117 | 0.999 | -0.013 | -16.546 | 42.409 | -0.390 | 0.999 | 0.675 |
| Left VIIIB | 3691 | -25.98 | 24.03 | -1.081 | 0.653 | -0.036 | -4.07 | 25.81 | -0.158 | 0.875 | -0.005 |
| Left IX | 3825 | -11.987 | 23.175 | -0.517 | 0.847 | -0.017 | 5.101 | 24.555 | 0.208 | 0.974 | 0.007 |
| Right Crus I | 3895 | -79.724 | 70.342 | -1.133 | 0.899 | -0.036 | 3.296 | 74.737 | 0.044 | 0.965 | 0.001 |
| Right Crus II | 3951 | -45.55 | 49.86 | -0.914 | 0.842 | -0.029 | -64.97 | 52.94 | -1.227 | 0.770 | -0.040 |
| Right VI | 3964 | 32.22 | 49.42 | 0.652 | 0.899 | 0.021 | -14.28 | 52.55 | -0.272 | 0.999 | -0.009 |
| **Right VIIB** | 3816 | **-121.60** | **41.84** | **-2.907** | **0.021*** | **-0.094** | 8.85 | 44.95 | 0.197 | 0.985 | 0.006 |
| Right VIIIA | 3604 | 7.788 | 35.990 | 0.216 | 0.967 | 0.007 | -82.665 | 38.834 | -2.129 | 0.231 | -0.072 |
| Right VIIIB | 3646 | 0.386 | 24.487 | 0.016 | 0.987 | 0.005 | -19.257 | 26.314 | -0.732 | 0.999 | -0.024 |
| Right IX | 3832 | -7.182 | 23.582 | -0.305 | 0.999 | -0.011 | -8.837 | 25.013 | -0.353 | 0.999 | -0.013 |
| **Flocculonodular** | | | | | | | | | | | |
| Left X | 3964 | -4.311 | 3.217 | -1.340 | 0.360 | -0.043 | -2.072 | 3.420 | -0.606 | 0.545 | -0.019 |
| **Right X** | 3961 | 0.7385 | 3.302 | 0.224 | 0.823 | 0.007 | **-8.282** | **3.515** | **-2.356** | **0.038*** | **-0.075** |
| **Vermis** | | | | | | | | | | | |
| **Vermis VI** | 3974 | **-24.015** | **8.705** | **-2.759** | **0.025*** | **-0.089** | 1.234 | 9.232 | 0.134 | 0.999 | 0.004 |
| Vermis VII | 3975 | -1.828 | 6.423 | -0.285 | 0.776 | -0.009 | -9.217 | 6.812 | -1.353 | 0.440 | -0.044 |
| Vermis VIII | 3977 | -19.966 | 11.876 | -1.681 | 0.233 | -0.054 | -14.073 | 12.635 | -1.114 | 0.442 | -0.036 |
| Vermis IX | 3973 | -3.994 | 11.866 | -0.337 | 0.920 | -0.012 | -26.372 | 12.503 | -2.109 | 0.175 | -0.087 |
| Vermis X | 3962 | -1.973 | 2.180 | -0.905 | 0.608 | -0.030 | 0.164 | 2.311 | 0.071 | 0.943 | 0.002 |
| Total Volume | 3978 | -677.4 | 394.4 | -1.718 | 0.086+ | -0.055 | 419.8 | 3964.7 | -0.839 | 0.401 | -0.027 |
| **Corpus Medullare** | 3948 | -40.583 | 78.276 | -0.518 | 0.604 | -0.017 | **-209.303** | **83.298** | **-2.513** | **0.012*** | **-0.080** |

Results of linear mixed effects models predicting cerebellar volumes including fixed effects of age, gender, PTSD diagnosis, major depressive disorder (MDD) diagnosis, intracranial volume and a fixed effect of site.

*$p < 0.05$, +$p < 0.10$

**Table S8:** *Effects of PTSD and AUD diagnosis on cerebellar volumes.*

| | | PTSD | | | | | AUD | | | | |
|---|---|---|---|---|---|---|---|---|---|---|---|
| **ROI** | **N** | ***b*** | ***SE*** | ***t*** | ***p*-FDR** | ***d*** | ***b*** | ***SE*** | ***t*** | ***p*-FDR** | ***d*** |
| **Anterior** | | | | | | | | | | | |
| Left I-III | 1225 | -10.600 | 12.530 | -0.846 | 0.999 | -0.051 | -32.131 | 17.934 | -1.792 | 0.222 | -0.111 |
| Left IV | 1229 | -13.032 | 34.860 | -0.374 | 0.709 | -0.021 | 41.260 | 50.009 | 0.825 | 0.409 | 0.047 |
| Left V | 1229 | 22.34 | 31.99 | 0.698 | 0.728 | 0.041 | -62.47 | 45.79 | -1.364 | 0.258 | -0.084 |
| Right I-III | 1228 | 0.499 | 13.409 | 0.037 | 0.970 | 0.002 | -1.819 | 19.207 | -0.095 | 0.999 | -0.006 |
| Right IV | 1220 | -3.76 | 36.78 | -0.102 | 0.999 | -0.006 | -16.76 | 52.81 | -0.317 | 0.999 | -0.018 |
| Right V | 1228 | -41.359 | 32.568 | -1.270 | 0.615 | -0.098 | -31.429 | 46.053 | -0.682 | 0.925 | -0.062 |
| **Posterior** | | | | | | | | | | | |
| Left Crus I | 1219 | -25.981 | 126.286 | -0.206 | 0.977 | -0.012 | -25.711 | 183.444 | -0.140 | 0.889 | -0.008 |
| Left Crus II | 1226 | 12.45 | 84.71 | 0.147 | 0.883 | 0.009 | -72.72 | 121.63 | -0.598 | 0.963 | -0.035 |
| Left VI | 1229 | -43.07 | 85.91 | -0.501 | 0.862 | -0.030 | -64.85 | 123.67 | -0.524 | 0.840 | -0.032 |
| Left VIIB | 1230 | -98.88 | 72.83 | -1.358 | 0.408 | -0.079 | 74.22 | 104.57 | 0.710 | 0.999 | 0.042 |
| Left VIIIA | 1230 | -125.67 | 70.68 | -1.778 | 0.266 | -0.103 | -170.47 | 101.44 | -1.680 | 0.651 | -0.099 |
| Left VIIIB | 1193 | -93.80 | 42.83 | -2.190 | 0.203 | -0.141 | -30.70 | 60.86 | -0.504 | 0.716 | -0.036 |
| Left IX | 1213 | 52.65 | 41.94 | 1.256 | 0.368 | 0.084 | -72.87 | 59.61 | -1.222 | 0.777 | -0.092 |
| Right Crus I | 1226 | -27.55 | 126.67 | -0.218 | 0.828 | -0.013 | -173.86 | 182.43 | -0.953 | 0.796 | -0.056 |
| Right Crus II | 1224 | -50.13 | 91.08 | -0.550 | 0.999 | -0.034 | 82.46 | 130.94 | 0.630 | 0.923 | 0.041 |
| Right VI | 1228 | 90.59 | 82.82 | 1.094 | 0.959 | 0.065 | -57.53 | 119.22 | -0.483 | 0.735 | -0.029 |
| Right VIIB | 1232 | -146.40 | 75.97 | -1.927 | 0.378 | -0.110 | -129.87 | 108.94 | -1.192 | 0.816 | -0.069 |
| Right VIIIA | 1213 | -42.780 | 61.996 | -0.692 | 0.999 | -0.045 | -51.360 | 87.880 | -0.584 | 0.783 | -0.042 |
| Right VIIIB | 1206 | -21.69 | 43.87 | -0.494 | 0.869 | -0.029 | -132.56 | 62.92 | -2.107 | 0.245 | -0.123 |
| Right IX | 1209 | 15.99 | 42.53 | 0.376 | 0.825 | 0.027 | -19.24 | 59.98 | -0.321 | 0.748 | -0.027 |
| **Flocculonodular** | | | | | | | | | | | |
| Left X | 1228 | -5.793 | 5.363 | -1.080 | 0.560 | -0.064 | -3.328 | 7.643 | -0.435 | 0.663 | -0.026 |
| **Right X** | 1234 | -5.256 | 5.598 | -0.939 | 0.348 | -0.054 | -5.236 | 8.022 | -0.653 | 0.514 | -0.037 |
| **Vermis** | | | | | | | | | | | |
| **Vermis VI** | 1231 | **-39.913** | **14.951** | **-2.670** | **0.035*** | **-0.163** | -9.577 | 21.408 | -0.447 | 0.819 | -0.029 |
| Vermis VII | 1229 | -3.111 | 11.604 | -0.268 | 0.789 | -0.016 | -2.780 | 16.547 | -0.168 | 0.867 | -0.011 |
| Vermis VIII | 1230 | -35.386 | 20.686 | -1.711 | 0.145 | -0.100 | -35.483 | 29.715 | -1.194 | 0.999 | -0.072 |
| Vermis IX | 1231 | -9.304 | 10.116 | -0.920 | 0.448 | -0.055 | -18.960 | 14.509 | -1.307 | 0.600 | -0.080 |
| Vermis X | 1232 | -7.848 | 3.996 | -1.964 | 0.125 | -0.122 | -6.281 | 5.688 | -1.104 | 0.675 | -0.072 |
| Total Volume | 1233 | -1028.72 | 715.46 | -1.438 | 0.151 | -0.083 | -1129.16 | 1029.80 | -1.096 | 0.273 | -0.063 |
| Corpus Medullare | 1227 | -240.35 | 126.26 | -1.904 | 0.057+ | -0.109 | -163.77 | 181.83 | 0.901 | 0.368 | -0.052 |

Results of linear mixed effects models predicting cerebellar volumes including fixed effects of age, gender, PTSD diagnosis, alcohol use disorder (AUD) diagnosis, intracranial volume, and a fixed effect of site.

$^{*}p < 0.05$, $^{+}p < 0.10$

**Table S9:** *Effects of PTSD and childhood trauma on cerebellar volumes.*

| ROI | N | PTSD *b* | PTSD *SE* | PTSD *t* | PTSD *p*-FDR | PTSD *d* | Childhood Trauma (CTQ) *b* | CTQ *SE* | CTQ *t* | CTQ *p*-FDR | CTQ *d* |
|---|---|---|---|---|---|---|---|---|---|---|---|
| **Anterior** | | | | | | | | | | | |
| Left I-III | 1132 | 1.883 | 15.431 | 0.122 | 0.903 | 0.007 | -0.551 | 7.935 | -0.069 | 0.945 | -0.004 |
| Left IV | 1124 | -47.17 | 38.28 | -1.232 | 0.327 | -0.074 | 14.11 | 19.65 | 0.718 | 0.710 | 0.043 |
| Left V | 1128 | 48.21 | 36.00 | 1.339 | 0.543 | 0.080 | -18.46 | 18.55 | -0.995 | 0.960 | -0.060 |
| Right I-III | 1135 | 13.637 | 16.153 | 0.844 | 0.600 | 0.050 | -2.324 | 8.310 | -0.280 | 0.999 | -0.017 |
| Right IV | 1122 | -19.43 | 40.96 | -0.474 | 0.635 | -0.028 | 17.91 | 21.05 | 0.851 | 0.395 | 0.051 |
| Right V | 1131 | -50.98 | 35.63 | -1.431 | 0.459 | -0.086 | 12.33 | 18.25 | 0.676 | 0.999 | 0.042 |
| **Posterior** | | | | | | | | | | | |
| Left Crus I | 1121 | -52.23 | 134.14 | -0.389 | 0.697 | -0.024 | -38.61 | 68.47 | -0.564 | 0.802 | -0.037 |
| Left Crus II | 1134 | -112.20 | 87.75 | -1.279 | 0.469 | -0.089 | 96.52 | 44.56 | 2.166 | 0.217 | 0.175 |
| Left VI | 1129 | -43.793 | 86.261 | -0.508 | 0.857 | -0.031 | 35.961 | 44.077 | 0.816 | 0.968 | 0.054 |
| Left VIIB | 1135 | -181.676 | 77.103 | -2.356 | 0.133 | -0.142 | 6.111 | 39.464 | 0.155 | 0.877 | 0.010 |
| Left VIIIA | 1125 | 35.091 | 74.717 | 0.470 | 0.746 | 0.028 | -42.430 | 38.422 | -1.104 | 0.945 | -0.067 |
| Left VIIIB | 1063 | -89.476 | 46.842 | -1.910 | 0.196 | -0.118 | -8.319 | 23.744 | -0.350 | 0.847 | -0.022 |
| Left IX | 1108 | -46.19 | 44.87 | -1.029 | 0.532 | -0.068 | 17.90 | 22.81 | 0.785 | 0.758 | 0.059 |
| Right Crus I | 1124 | -108.711 | 131.795 | -0.825 | 0.574 | -0.053 | -2.565 | 66.902 | -0.038 | 0.969 | -0.003 |
| Right Crus II | 1133 | -126.42 | 91.17 | -1.387 | 0.291 | -0.092 | -19.24 | 46.23 | -0.416 | 0.948 | -0.031 |
| Right VI | 1131 | -3.80 | 84.94 | -0.045 | 0.964 | -0.003 | 53.32 | 43.31 | 1.231 | 0.383 | 0.087 |
| Right VIIB | 1137 | -153.25 | 84.93 | -1.804 | 0.497 | -0.108 | 62.00 | 43.63 | 1.421 | 0.999 | 0.086 |
| Right VIIIA | 1113 | 23.04 | 62.31 | 0.370 | 0.831 | 0.024 | -40.51 | 31.38 | -1.291 | 0.690 | -0.092 |
| Right VIIIB | 1092 | -70.206 | 48.640 | -1.443 | 0.348 | -0.088 | 4.574 | 24.777 | 0.185 | 0.996 | 0.011 |
| Right IX | 1104 | -81.87 | 46.81 | -1.749 | 0.284 | -0.119 | 29.64 | 23.74 | 1.248 | 0.495 | 0.096 |
| **Flocculonodular** | | | | | | | | | | | |
| Left X | 1137 | -10.810 | 6.130 | -1.763 | 0.156 | -0.109 | 0.235 | 3.135 | 0.075 | 0.940 | 0.005 |
| Right X | 1137 | -7.689 | 6.328 | -1.215 | 0.225 | -0.072 | -0.514 | 3.251 | -0.158 | 0.874 | -0.009 |
| **Vermis** | | | | | | | | | | | |
| Vermis VI | 1135 | -40.726 | 16.719 | -2.436 | 0.075+ | -0.152 | 2.155 | 8.540 | 0.252 | 0.999 | 0.017 |
| Vermis VII | 1137 | -3.892 | 12.465 | -0.312 | 0.755 | -0.019 | -9.323 | 6.358 | -1.467 | 0.715 | -0.095 |
| Vermis VIII | 1134 | -34.874 | 21.776 | -1.602 | 0.275 | -0.098 | 7.222 | 11.162 | 0.647 | 0.863 | 0.042 |
| Vermis IX | 1137 | -4.814 | 10.679 | -0.451 | 0.815 | -0.029 | -0.156 | 5.442 | -0.029 | 0.977 | -0.002 |
| Vermis X | 1133 | 3.505 | 4.258 | 0.823 | 0.685 | 0.052 | -2.354 | 2.173 | -1.083 | 0.698 | -0.075 |
| Total Volume | 1141 | -1216.956 | 719.835 | -1.691 | 0.091+ | -0.103 | 3.753 | 368.084 | 0.010 | 0.995 | 0.001 |
| Corpus Medullare | 1134 | -125.87 | 151.99 | -0.828 | 0.408 | -0.049 | -41.35 | 78.22 | -0.529 | 0.597 | -0.031 |

Results of linear mixed effects models predicting cerebellar volumes including fixed effects of age, gender, PTSD diagnosis, childhood trauma severity (CTQ), intracranial volume, and a fixed effect of site.

+$p < 0.10$
